# Dynamic neurogenomic responses to social interactions and dominance outcomes in female paper wasps

**DOI:** 10.1101/2021.03.01.433260

**Authors:** Floria M.K. Uy, Christopher M. Jernigan, Natalie C. Zaba, Eshan Mehrotra, Sara E. Miller, Michael J. Sheehan

**Author notes:** Present address: Department of Biology, University of Rochester, Rochester, NY 14627.

## Abstract

Social interactions have large effects on individual physiology and fitness. In the immediate sense, social stimuli are often highly salient and engaging. Over longer time scales, competitive interactions often lead to distinct social ranks and differences in physiology and behavior. Understanding how initial responses lead to longer-term effects of social interactions requires examining the changes in responses over time. Here we examined the effects of social interactions on transcriptomic signatures at two points, at the end of a 45-minute interaction and 4 hours later, in female *Polistes fuscatus* paper wasp foundresses. Female *P. fuscatus* have variable facial patterns that are used for visual individual recognition, so we separately examined the transcriptional dynamics in the optic lobe and the central brain. Results demonstrate much stronger transcriptional responses to social interactions in the central brain compared to the optic lobe. Differentially regulated genes in response to social interactions are enriched for memory-related transcripts. Comparisons between winners and losers of the encounters revealed similar overall transcriptional profiles at the end of an interaction, which significantly diverged over the course of 4 hours, with losers showing changes in expression levels of genes associated with aggression and reproduction in paper wasps. On nests, subordinate foundresses are less aggressive, do more foraging and lay fewer eggs compared to dominant foundresses and we find losers shift expression of many genes, including vitellogenin, related to aggression, worker behavior, and reproduction within hours of losing an encounter. These results highlight the early neurogenomic changes that likely contribute to behavioral and physiological effects of social status changes in a social insect.

## INTRODUCTION

Social interactions can give rise to a range of immediate as well as long-lasting effects on behavior and physiology^1–4^. Regardless of the nature of the interaction or the outcome, social experiences are expected to have a number of shared effects on the physiology of those involved. Processing social information may depend on multiple cues or signals, which may be processed by generalized or social-specific cognitive mechanisms^5^. In addition to social information processing, interactions can increase rates of activity and movement, especially in relation to courting or fighting^2,6^. Longer-term consequences of social interactions depend on the nature and outcome of the encounters. Cooperative interactions can lead to benefits for multiple individuals as well as physiological responses that aid in reinforcing social bonds. Competitive interactions, in contrast, often lead to divergent outcomes for individuals –i.e., a winner and loser. Winning versus losing typically cause different physiological and behavioral responses^7–13^. Over repeated interactions, this can lead to profound differences in behavior, physiology, life expectancy, and fitness^4,14–17^.

How are social outcomes translated into physiological changes? Ultimately, the answer to this question lies at the intersection of the neural circuits that process information as well as the resulting neurogenomic shifts, i.e., the changes in patterns of brain gene expression, that accompany social challenges. In recent years there has been a growing number of gene expression studies examining the neurogenomic responses to social interactions across a range of taxa including honeybees, mice and sticklebacks^6,18,19^. In a broad sense, social interactions are expected to engage similar brain circuits across individuals. For example, in vertebrates these brain regions have largely been conserved across 450 million years of evolution ^20^. Indeed, at the level of neural firing patterns, social interactions give rise to similar patterns of neural activity in bats and mice^21,22^. While a similar network has not been identified across insects, we might reasonably expect members of the same species to engage similar brain regions and likely have similar initial neurogenomic responses to social interactions as well.

Divergent social outcomes lead to different physiological responses, which may be initiated by differences in neurogenomic responses shortly following an interaction. There have also been studies examining the effects of winning and losing rather than simply the response to challenge *perse*. In zebrafish, socially driven transcriptional changes require individuals to assess the outcome of the interaction^23^ (i.e., did they win or lose). In sub-social carpenter bees, repeatedly winning or losing staged contests gives rise to distinct neurogenomic profiles^11,24^. In the ant *Harpegnathos saltator*, workers compete for reproductive openings upon the removal of the queen and within a few days individuals have divergent neurogenomic profiles depending on their trajectory toward either staying as a worker or becoming a reproductive gamergate^25^. Similar divergence in social behavior and neurogenomic profiles are seen among *Polistes dominula* paper wasp workers upon queen removal^26,27^. Collectively, these studies demonstrate that social interactions can have immediate effects and that repeated interactions can have longer-term consequences for patterns of transcription in the brain that differ for winners and losers or higher-versus lower-ranking individuals. Understanding how transcriptional patterns change over time in response to different social interactions and across different taxa will help us to more clearly link social outcomes to short and long-term physiological changes.

Understanding the dynamic changes that occur between initial responses and subsequent divergence between winners and losers will help link these two areas of research. Studies examining the temporal dynamics of transcriptional responses to social challenge in stickleback and mice over the course of a few hours^18,19^ highlight the transient and dynamic nature of transcriptional responses. Detailed work on the early transcriptional responses to fighting between pairs of male beta fish demonstrates that fighting individuals have shared transcriptomic responses within the first hour after fighting^28^. Though the studies mentioned above have looked at dynamic responses to a social challenge from territorial or nest intrusions or more established winner-loser effects, the dynamics by which interacting individuals develop divergent transcriptomic responses over the course of a few hours has received less attention.

Here we examine the dynamic neurogenomic responses to social interactions in female *Polistes fuscatus* paper wasp foundresses over the course of four hours following a staged social interaction. Paper wasps are primitively eusocial insects in which females found new nests each spring after overwintering^29^. Social interactions among paper wasp foundresses lead to profound physiological differences between dominants and subordinates. Nests are initiated by a single foundress or small groups of foundresses, who form an aggression-based dominance hierarchy, which determines the extent of work and egg-laying^30,31^. Polistine foundresses have aggressive interactions in both the pre-nesting stage as well as on the nests, where they interact aggressively with co-foundresses as well as occasional usurpers^32–35^. Wasps also reliably show aggression to other individuals in neutral arenas, providing a convenient method for studying the effects of aggression in a controlled setting^36–38^. Previous work has shown that *Polistes* foundresses respond rapidly to aggressive encounters by modulating juvenile hormone^13^, though genome-wide transcriptomic responses have yet to be examined immediately following aggressive interactions. In established co-foundress associations, dominant and subordinate foundresses show differential expression of genes associated with aggressive behavior^39^. By comparing the temporal shifts in gene expression between winners and losers, we can potentially identify genes that are associated with the early stages of dominance hierarchy formation in paper wasps, as well as generate more general insights into the neurogenomic processes by which social interactions lead to divergence in behavior and physiology.

The neurogenomic responses to social interactions in *P. fuscatus* are also of interest because this species recognizes individuals based on variable facial features^5,40^. Individual recognition appears to mediate dominance interactions among groups in the lab and on natural nests^37,40^. Individual recognition is not present in other closely related species of paper wasps^5,41^, suggesting the trait has evolved relatively recently^42^. Neurogenomic responses to operant conditioning related to face-learning have been previously studied^43^, but their neurogenomic responses to social interactions have not been investigated. Wasps are known to form long-term memories of those they have interacted with^44^, so examination of neural transcriptomes a few hours after the interaction has the potential to reveal insights into the neurogenomic responses related to social memory, as long-term memory formation occurs hours after initial learning has occurred^45^. Given the importance of vision in social interactions for this species, we examined the effects of social interaction on the optic lobe as well as the central brain (Fig 1a, hereafter ‘optic lobe’ and ‘brain’).

**Figure 1:**
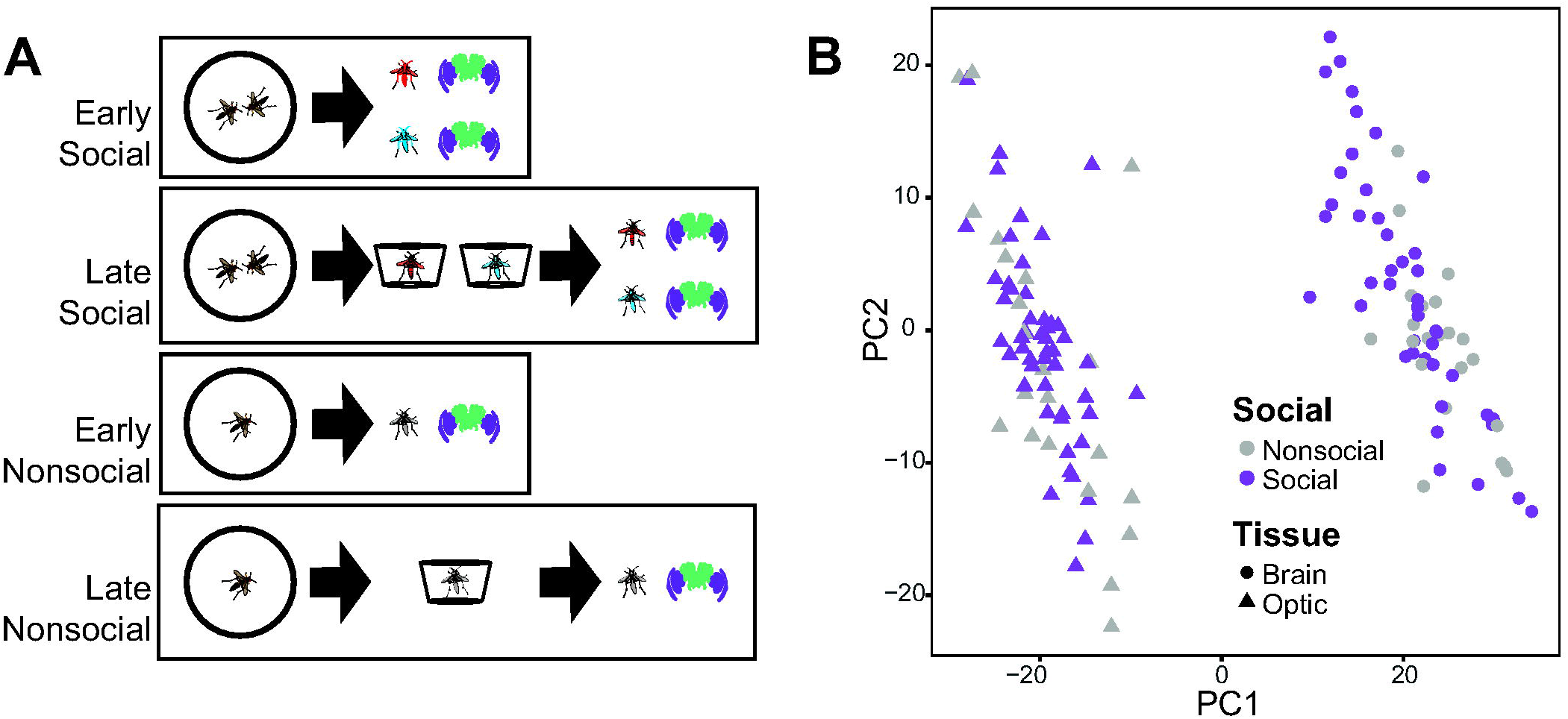
Overview of experimental design and RNAseq data. (A) The experiment consisted of generating two groups of wild-caught wasps that either engaged in a recent social experience or remained nonsocial. Half of each group was sacrificed at the end of a 45-minute interaction period with the other half held in individual containers for 4 hours until they were then sacrificed. RNA was extracted separately from the combined optic lobes (purple) and the remainder of the brain, called ‘brain’ throughout (green). In other figures, we show the part the tissue the data is derived from with the relevant icon. (B) Tissue is the strongest separator of the data in a principal component analysis. Within the brain, but not the optic lobe, social experience also has a major influence on neurogenomic patterns. Here and in subsequent figures, red wasp symbols are used to indicate winners, blue wasp symbols for losers, and grey wasps for control individuals that did not have social interactions.

We designed an experiment to examine the dynamic neurogenomic responses shortly after social interactions in the optic lobe and non-visual brain (Fig 1a). Wasps were filmed in a neutral arena while paired with another weight-matched individual or alone. To better understand the temporal dynamics of neurogenomic responses in the hours following a social interaction, we looked at transcriptomes at two time points: immediately following a 45-minute interaction and after 4 hours of separation back in the wasps’ original housing containers (Fig 1a). In the grander scheme of paper wasp dominance relationships, both of these timepoints are very early in the time course over which a dominance hierarchy would be formed. For ease of distinguishing between the samples we refer to those taken immediately at the end of a 45-minute interaction as ‘early’ and those at 4 hours as ‘late’.

Using the RNAseq data from paper wasp foundresses, we address multiple questions. (1) How does the magnitude of neurogenomic responses differ between peripheral and central processing? To the extent that responses are driven by the processing of social outcomes rather than simply response to social stimuli, we may expect larger and or more dynamic changes in more central compared to peripheral brain regions. (2) Given that paper wasps learn and remember the identities of wasps they interact with^44^, is there a detectable neurogenomic signature related to memory in paper wasps following interactions? (3) How does social outcome influence the dynamics of neurogenomic responses over the course of a few hours? Recent studies suggest similar neural responses among individual during or right after social interactions^21,22,28^, whereas others demonstrate divergent outcomes over the course of days^11,24,25,27^. Therefore, we may predict that initial neurogenomic responses will be more similar immediately following social interactions and that winners and losers will diverge transcriptionally over time.

## RESULTS AND DISCUSSION

### Social interactions generate stronger and more dynamic neurogenomic responses in the central brain compared to optic lobe

We first compared RNAseq data from 139 samples in DESeq2 with a model that included tissue (optic lobe or central brain), whether or not wasps had been placed in a social or control trial, and time of sacrifice as separate categorical main factors. Optic lobes and the brain show distinct transcriptional profiles that are well-separated in a PCA (Fig 1c). We identified 4937 differentially expressed genes (DEGs) between the brain and optic lobes consistent with different cellular compositions between the two tissues. Time of sacrifice showed a minor effect on overall patterns of gene expression with 73 DEGs. In contrast, social experience had a more pronounced effect on patterns of gene expression, with 742 DEGs (Fig S1). Furthermore, social and non-social samples are better separated in principal component space among brain samples compared to optic lobe (Fig 1b). Though social and nonsocial central brain samples are differentiated along PC2 (ANOVA, *F*_164_ = 4.75, *P* = 0.033), the groups do not form two distinct clusters as has been found in other transcriptomic studies related to social behavior in other species (e.g. Vu et al. 2020). The behavioral paradigm used in this study mirrors other lab studies of social behavior and cognition in *P. fuscatus* that examined encounters in a neutral arena and detect variable amounts of aggression^41,44,46^, though is likely to be a less extreme social experience compared to paradigms that challenge individuals in their nest or home cage and or otherwise produce strong fighting responses used in other behavioral transcriptomic studies^6,18,19,28^. Although the social experiences in our trials were comparatively mild, we nevertheless detect hundreds of differentially expressed genes in response to social interactions.

We next considered a model comparing each group based on brain region, time and social experience as a single combined factor (e.g., brain_early_social v. brain_early_nonsocial). Consistent with visual separation in the PCA (Fig 1b), the comparisons reveal a stronger effect of social interactions on the brain compared to the optic lobe (Fig 2a). The results are qualitatively similar when examining the effects of social experience and time on brain and optic datasets separately (Fig 2b-c, Table S2).

**Figure 2:**
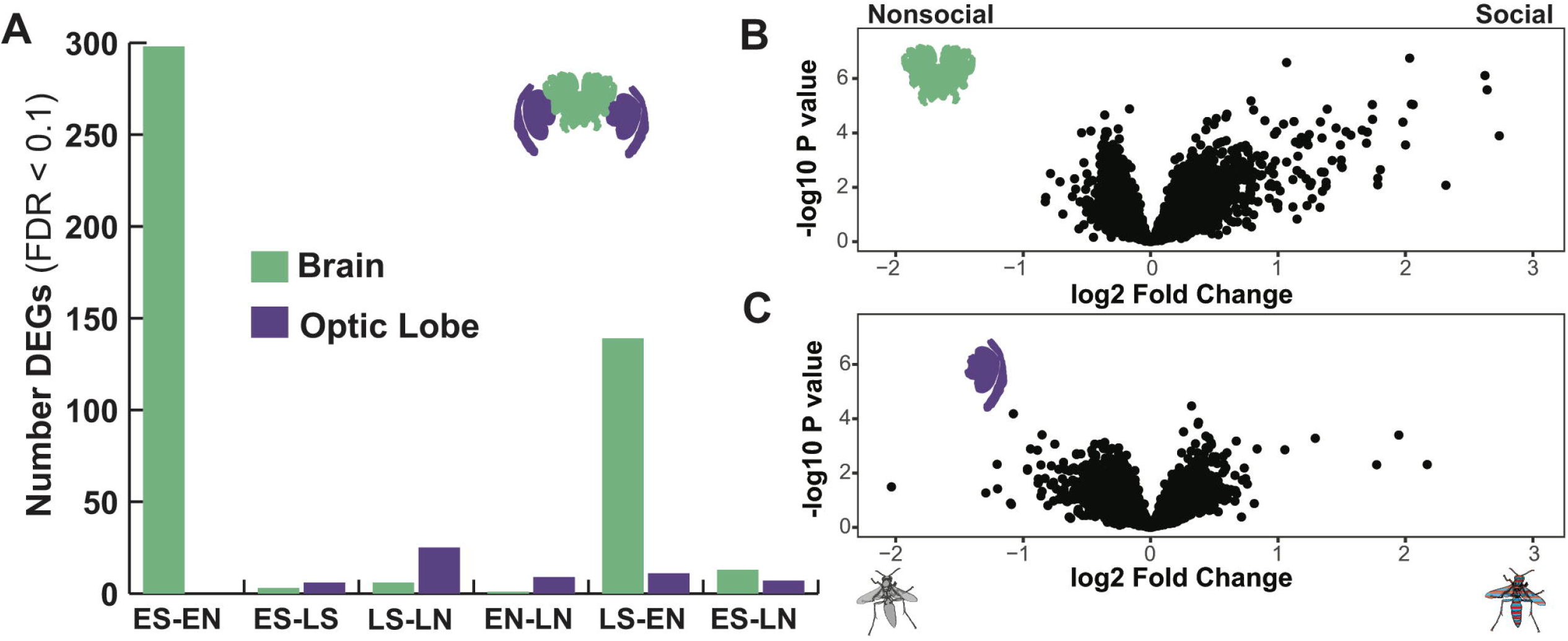
Social interactions influence neurogenomic signatures more in the brain than optic lobe. (A) The effects of social interactions are stronger in the brain compared the optic lobe. At both early and late time points there are hundreds of genes differentially expressed (FDR < 0.1) between social and nonsocial groups. The following codes are used in the axis legend: ES = early social, EN = early nonsocial, LS = late social, LN = late nonsocial. (B) The volcano plots show the log2 fold change between social (up) and nonsocial (down) on the x-axis and the – log10 P value. The red and blue striped wasp symbol indicates that the data includes all socially interacting wasps.

These data add to a growing body of literature documenting the changes in brain transcriptomic profiles in response to social behavior^2,6,11,19,24,28^. Consistent with those studies, we find hundreds of genes that are differentially regulated in some comparisons. The neurogenomic effects of social interaction are detectable at both the earlier (at the end of a 45-minute interaction) and later (4 hours following the interaction) time points, but the evidence for differential gene expression between social and nonsocial individuals is strongest shortly following an interaction (Fig 2a). The transcriptomic signatures measured right after the interaction represent a combination of immediate responses to social stimuli and interactions as well as some of the initial downstream physiological responses to social behavior. In contrast, at the 4 hour timepoint individuals had been removed from social interactions for a period of time so socially regulated genes at this later timepoint reflect downstream consequence of social interactions^3^. The increased number of differentially expressed genes at the earlier timepoint may reflect the engagement of a broad set of neural circuits and gene-networks during social interactions. Conversely, the decrease in differential expression over time could also reflect divergent response to social outcomes from winners and losers, such that there is more ‘noise’ in the transcriptomic signatures of the wasps with recent social experience after a few hours (see below for follow up analyses).

There is a growing literature demonstrating that sensory system tuning and function is more dynamic and plastic than has been previously appreciated^47–51^. Though examples of sensory plasticity are often developmental shifts in response to predictable cues such as season or reproductive state, there is also evidence that individuals’ sensory systems respond to their physical environment^52^. We examined the responses of optic lobes to social interactions in paper wasps and found modest evidence of differential expression 4 hours after social interactions (Fig 2a). Among the differentially expressed genes include a dopamine transporter and a major royal jelly protein, which are both downregulated in the 4-hour time point in social compared to nonsocial wasps, suggesting the possibility for modulatory effects on the visual system following social interactions. It is possible that longer-term exposure to social interaction or isolation could have more dramatic effects on visual systems. Indeed, social experience during development is required for individual recognition in *P. fuscatus^53^*.

### Socially responsive genes are enriched for memory-related functions

We identified 61 overrepresented GO terms (P < 0.01) among the 742 social DEGs in the full model with brain region, social experience, and sampling time as separate categorical factors. Many of the GO terms deal with membrane transport, calcium signaling, synaptic transmission or behaviors, which are to be expected given that we analyzed a neurogenomic dataset related to adult behavior (Table S3). A number of the enriched categories, however, suggest other neurogenomic processes supporting social behavior in *Polistes* wasps. For example, genes annotated as being involved in cholinergic synaptic transmission are overrepresented among socially responsive genes (GO:0007271, P = 0.0015), suggesting that cholinergic neurons may play a role in the aggressive encounters between the wasps. Recent work in *Drosophila* has implicated cholinergic signaling in aggression in both males and females^54–56^, suggesting potentially shared mechanisms related to aggressive interactions across taxa.

Female *P. fuscatus* learn and remember the identity of other wasps from previous interactions^44^ or even outcomes of fights among other individuals they have seen interacting^46^. Behavioral experiments have demonstrated both short and long-term memories of individuals^44,46^, suggesting that signatures of both processes may be enriched among differentially regulated genes. Indeed, genes annotated with functions in anesthesia-resistant memory (GO:0007615, P = 3.6e-5) and long-term memory (GO:0007616, P = 0.009) are enriched among socially responsive genes. Anesthesia-resistant memory refers to a process of memory consolidation that is resistant to disruptions in neural activity, as would be caused by anesthesia^57^. It does not require protein synthesis and is considered a form of intermediate-term memory^58,59^. Long-term memory in contrast requires protein synthesis and the reweighting of synaptic connections^60,61^. A puzzling feature of the expression of genes annotated with memory functions is that they frequently appear to be down regulated among individuals in the social compared to nonsocial treatments (Fig S1). Memory formation is a dynamic process with multiple steps in which genes are up- and down-regulated at different times^62^ and the observed down-regulation may reflect aspects of that dynamics process. Most studies of the genetic basis of memory formation in invertebrates have focused on single cue associations (e.g., a color or smell) but the social interactions studied here are more complex in terms of sensory inputs and the range of positive and negative experiences that occur. Global downregulation in the brain may mask upregulation in specific neurons where social memories are encoded. While these data demonstrate that social interactions influence the expression of memory-related genes, understanding how these patterns translate to memory formation (or lack thereof) will require further study.

Likely relevant to memory formation, socially responsive genes are enriched for functions relating to mushroom body development (GO:0016319, P = 0.00055), synaptic target recognition (GO:008039, P = 0.00029), and regulation of synaptic plasticity (GO:0048167, P = 0.0051). Long-term memory formation requires modulation of synaptic connections^62^, which may be captured by GO terms dealing with changes to synapses including their plasticity and targeting. Additionally, enrichment for GO terms related to mushroom body development when seen in the context of an adult brain, are suggestive of a role of mushroom body neuropils in social processing and memory. The context or features of an interaction that make it more or less memorable for paper wasps remain to be investigated, though the present study was able to detect neurogenomic signatures related to memory following interactions in a neutral arena. How investment in memory may vary across social contexts (on a nest versus a neutral arena) and the intensity of the interactions are open questions that the present data suggest could be addressed, at least in part, using transcriptomic techniques.

### Similarities and differences in winner and loser neurogenomic responses

Individual wasps had different experiences of social interactions depending on whether or not they were the individual giving or receiving more aggression – i.e., whether they were the winner or the loser of the encounter. Therefore, we considered the neurogenomic responses separately for the individuals that won or lost the social encounters compared to those that had not been involved in a social interaction. In a model considering encounter outcome, tissue, and time as main factors, we found overall similar numbers of DEGs for tissue (4435 DEGs) and time (22 DEGs) as with the model based on social experience. Both winners and losers had hundreds of differentially expressed genes compared to nonsocial individuals, though the neurogenomic response appears to be stronger in losers (Fig 3a, winners = 217 DEGs, losers = 584 DEGs). When directly compared to each other, winners and losers show no significant differences in gene expression based on the FDR < 0.1 threshold in DESeq2. Even considering less restrictive criteria for calling DEGs, only 55 genes have P < 0.01 when not correcting for false discovery rates. The lack of strong differential expression between winners and losers suggests that the two social outcomes have similar expression profiles when analyzing the entire dataset, including both brain regions and timepoints. Indeed, there are 113 DEGs shared between winners and losers, a significantly greater overlap than expected by chance (Fig 3a, P < 2e-16). Both winners and losers also show significant overlap with the DEGs responding to social interactions in general (P < 2e-16 in both cases). Next, we compared the patterns of differential expression of winners and loser in relation to the nonsocial wasps. The log2 fold changes in both winners and losers compared to nonsocial wasps in the entire dataset are strongly correlated (Fig3b, linear model: y = 0.84x - 0.02, *F*_1,4002_ = 7458, *r^2^* = 0.65, P < 2e-16). Thus, when considering the entire dataset encompassing both brain regions and sampling points, winners and losers have broadly similar responses, though with a greater number of DEGs in losers compared to the nonsocial individuals (Fig 3).

**Figure 3:**
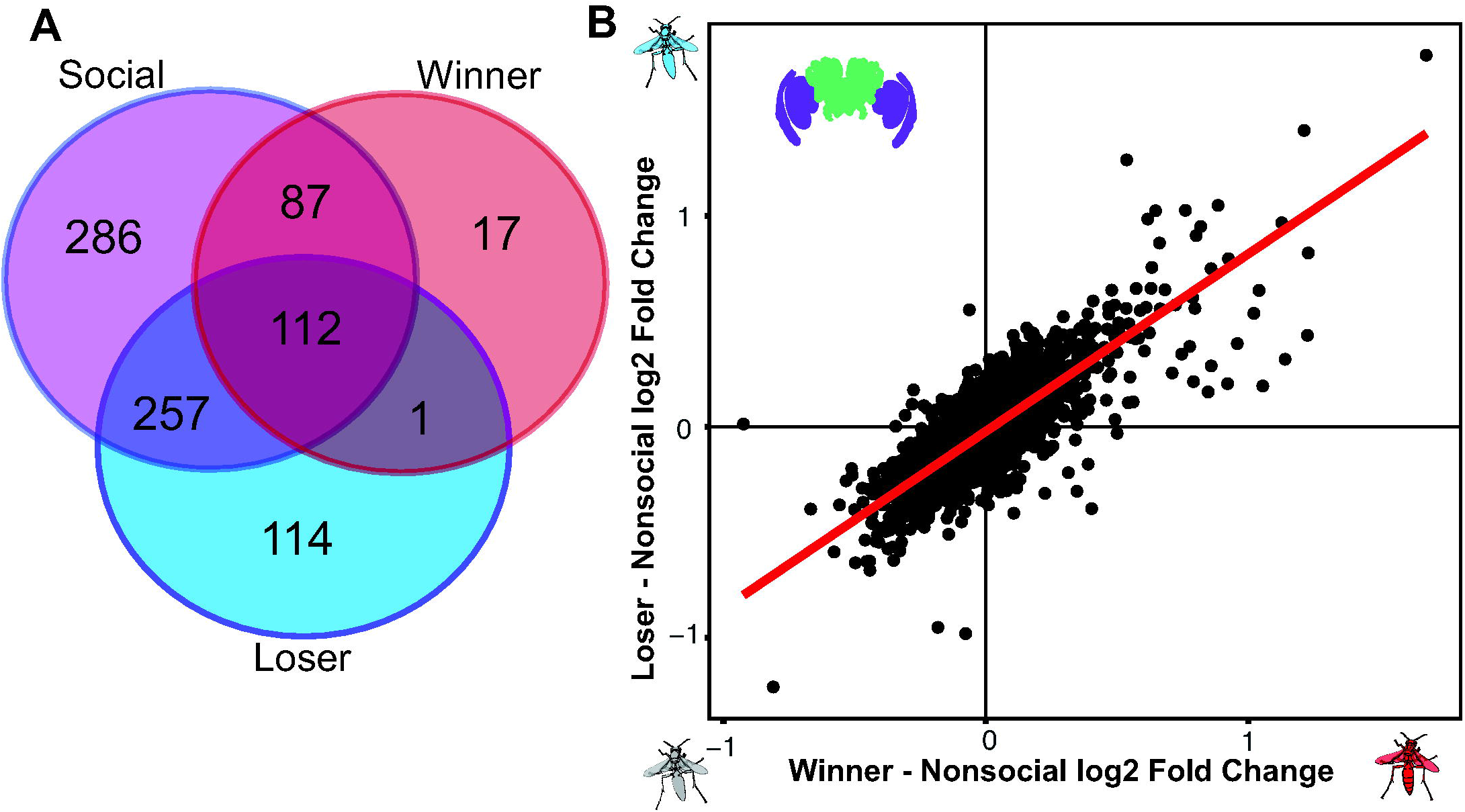
Similar overall neurogenomic responses in winners and losers. (A) There is significantly more overlap than expected by chance between the DEGs for winners and loser compared to each other as well as both winner and loser compared to all individuals with recent social experience (P < 2e-16). (B) The difference in log2 fold change in gene expression for all genes with a mean expression count of 100 or greater for nonsocial individuals are correlated for winners and losers. Both panels show analyses from the entire dataset with both brain regions and time points combined.

We investigated the relationship between gene expression patterns in winners and losers further by comparing the patterns of differential expression relative to nonsocial individuals at the end of the 45-minute interaction and 4 hours later. Here we present the results of gene expression in the non-visual brain since we observed stronger effects of social behavior in the brain than optic lobe (Table S4). We examined the log2 fold change in expression in losers relative to nonsocial individuals in a mixed model with winner log2 fold change relative to nonsocial individuals and time as fixed effects and gene as a random effect. Differential expression between winners and nonsocial wasps predicts expression differences in losers relative to nonsocial wasps (t = 69.02, df = 7420, P < 2e-16). Time was a significant predictor with greater log2 fold changes in losers compared to nonsocial wasps at the later time point (t = 12.27, df = 3313, P < 2e-16). There was a significant interaction between the extent of differential expression between winners and nonsocial wasps and time (t = 3.3, df = 5424, P = 0.00096). Next, we calculated a separate regression between loser and winner responses compared to nonsocial individuals at early and later times to further investigate these patterns. The slope of the regression is steeper though the fit substantially poorer between winners and losers at the later timepoint compared the earlier sampling time (Fig 4a, early: y=0.69x + 0.001, *r^2^* = 0.70; Fig 4b, later: y=0.74x - 0.06, *r^2^* = 0.38).

**Figure 4:**
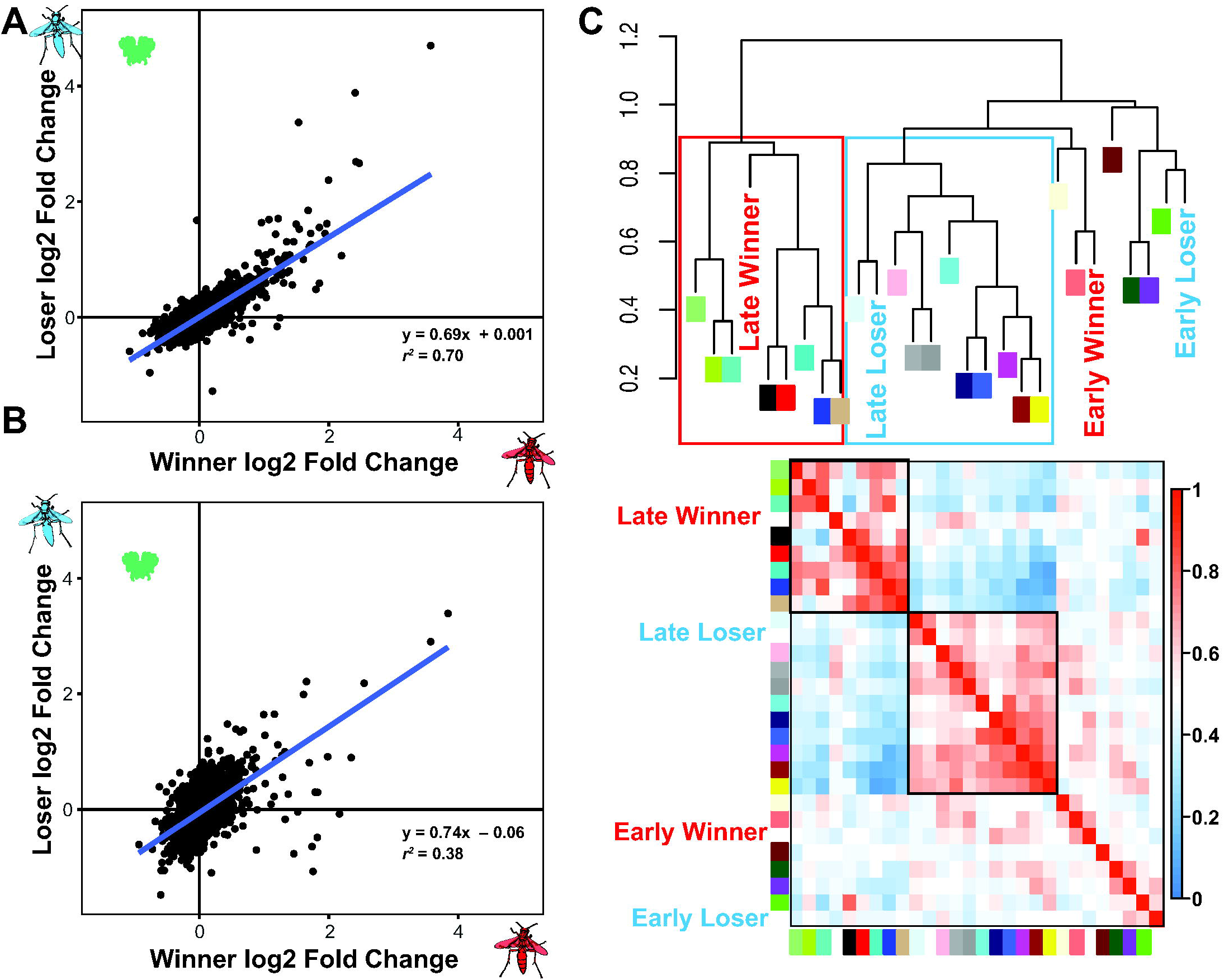
Divergence in loser brain transcriptomes over time. (A) Focusing on only the brain dataset, the log2 fold change in gene expression differences between nonsocial individuals and winners and losers are well correlated at the earlier time point. (B) At the later time point, there is substantially less correlation between winner and loser responses relative to nonsocial individuals. (C) Gene correlation modules are organized into two meta-modules, which are associated with late winners and late losers respectively. The top panel shows a dendrogram with the colors labeled and social outcomes labeled. The boxes have been added to highlight the two meta-modules. The bottom panel shows a heatmap showing the relationships among modules. Higher correlations are show by warmer red colors with modules with low or not correlations shown in blue. The two meta-modules highlighted in the dendrogram have been highlighted here with black outlines.

Winners and losers show a pattern of increased divergence in non-visual brain gene expression over time using a distinct analysis method as well. We used weighted correlation network analysis (WGCNA) to examine patterns of co-expressed genes in relation to social behavior^63^. WGCNA assigned 6086 genes to 24 modules (mean = 253.58 genes, max = 1091, min = 39). Multiple modules are significantly associated with winning or losing an encounter. Co-expression modules associated with winning or losing at either time point are all distinct – i.e., no modules are correlated with more than one outcome-time combination (Fig S2). We examined the relationship among modules and social behaviors by identifying meta-modules, correlated groups of eigengenes, and examining their relationship with different social outcomes. The brain dataset contains two large meta-modules that are associated with late winners and late loser respectively (Fig 4c). In contrast, early sampled losers and winners do not group within clear meta-modules. WGCNA calculates modules blind to the sample attributes such as time of sampling, whether wasps had been given a social experience, or the outcome of that interaction. Nevertheless, WGCNA identifies two distinct gene co-expression meta-modules associated with late-sampled losers and winners respectively reinforcing the observation that antagonistic social interactions lead to increased divergence in neurogenomic states over time.

Taken together, these data suggest that the overall neurogenomic responses to social interactions are similar in winners and losers observed in the whole dataset is driven by their initial similarity at the end of the interaction. The responses diverge over the course of a few hours, with relatively greater differences relative to individuals that did not experience social encounters appearing in losers over time. The correlation between winners and losers at the early time point echoes shared patterns of neural activity observed in mice and bats or shared transcriptomic signatures among interacting individuals in beta fish^21,22,28^. Given that competitive social interactions typically lead to divergent outcomes for winners and losers or dominants and subordinates^4,13,17,64,65^, the initial similarity in neural responses between competing individuals may seem counterintuitive. The similar neurogenomic responses of winners and losers observed at the earlier timepoint, however, declines over time in our dataset. The similar early responses may reflect the activity of neural mechanisms for assessing social stimuli and the initial processing of the encounter that is shared between the interacting individuals. Divergence over time may reflect the integration of the outcome into neurogenomic responses that themselves go on to further influence behavioral states following social encounters. This divergence among socially interacting wasps likely contributes to the reduced number of differentially expressed genes detected between social and nonsocial treatments at the late time point due to heterogeneity in expression patterns between winners and losers. Reproductive division of labor among groups of foundresses is based on physical aggression in *Polistes*^29,30^, but ultimately results in distinct neural and physiological states between the dominant and subordinate foundresses^39,66^. Understanding the steps that lead from similar to divergent neurogenomic states between interacting individuals will help clarify how social experiences come to generate diversity in physiology and behavior among individuals in a population ^3,67^.

### Dynamics changes in gene expression in the hours following a social interaction depends on dominance outcome

To investigate the neurogenomic changes that may accompany shifts associated with winning or losing, we compared the relative magnitude of brain gene expression changes between early and late losers to those seen between early and late winners (Fig 5). There is a statistically significant but very weak negative relationship between the relative changes seen in winners compared to losers (Fig 5: F_2,3709_=23.39, P = 1.55e-12, *r^2^* = 0.014). Consistent with the previous analyses (Fig 4), we find that there are more extreme changes in losers compared to winners, shown by the greater spread along the y-axis (Fig 5). Interestingly, this observation fits with theoretical results that loser-effects should be stronger than winner-effects^68^.

**Figure 5:**
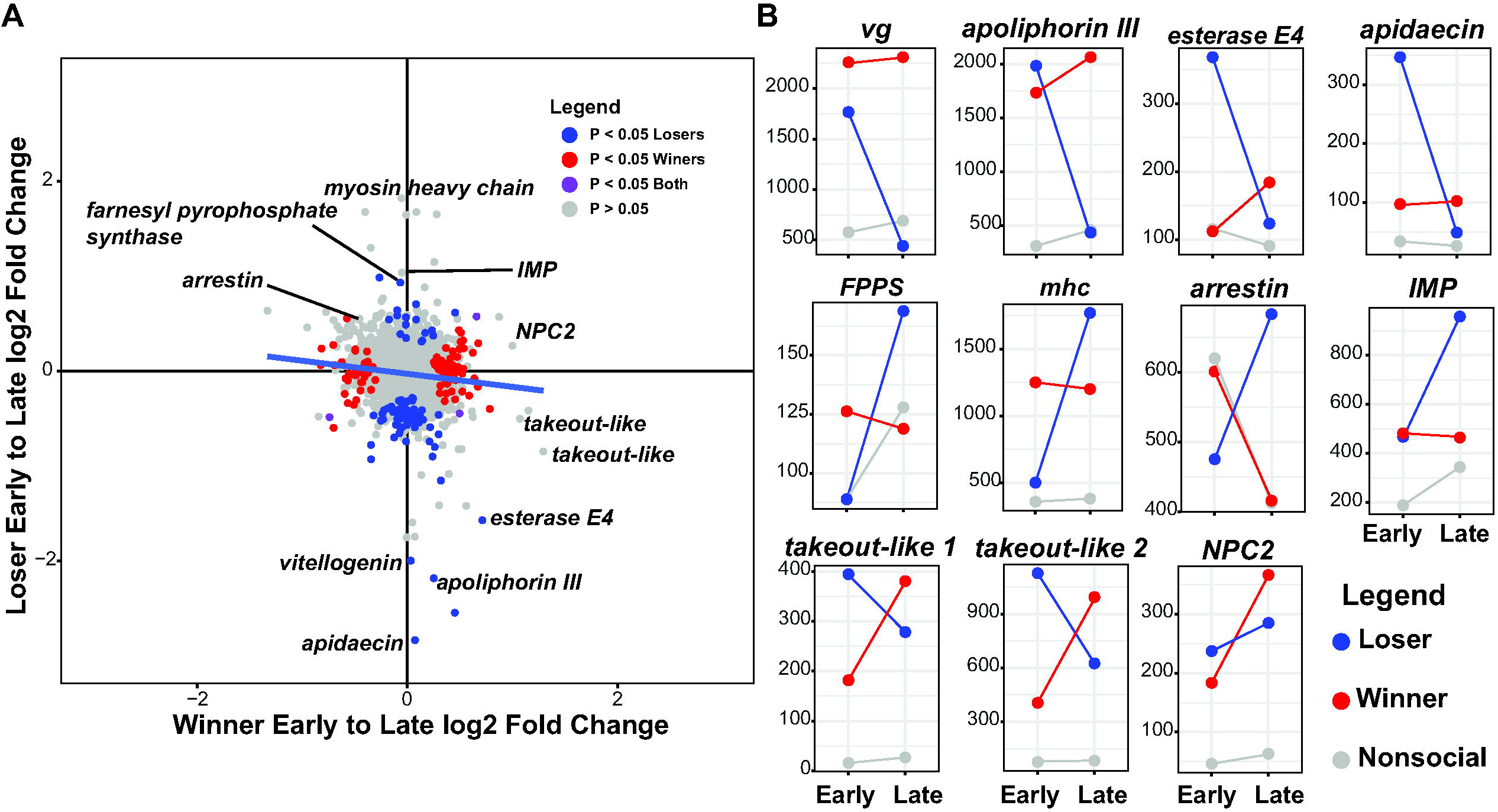
Shifts in winner and loser gene expression over time. (A) There are more dramatic shifts in the responses of losers compared to winners over time. The scatter plot shows the log2 fold change between early and late winners on the x-axis against the similar early to late comparison for losers on the y-axis. Thus, genes in the upper right quadrant are those that increase over time in both winners and losers, while those the upper left quadrant increase in losers but decrease in winners. The greater spread along the y-compared to x-axis shows that there are larger changes in loser gene expression profiles over time compared to winners. There is a weak but significant negative correlation suggesting that some genes that increase in losers tend to decrease in winner and vice versa. Notable gene are highlighted. Data points are color-coded according to the legend. (B) The panels show the mean normalized count of expression for losers, winners and nonsocial individuals at early and late sampling points. Lines are drawn connecting the points between groups of the same social outcome. Note that the y-axis is different for each gene and depends on the dynamic range of the specific gene. For example, *arrestin* shows a much smaller change in expression across groups than *takeout-like 1*, which is expressed at very low levels in nonsocial controls but expressed much more highly in wasps that engaged in social interactions.

We next examined the identity of genes with extreme changes in both winners and losers to learn more about the nature of neurogenomic changes. Notable genes are highlighted in Fig 5. We observe multiple patterns of change including genes that are initially upregulated in losers relative to nonsocial wasps at the early time point and then substantially decreased at the later time point. Many of the genes with the largest decreases in losers at the later time show this up-then-down pattern, including *vitellogenin, apolipophorin III, esterase E4* and *apideacin*. Both *vitellogenin* and *esterase E4* are consistently downregulated in workers compared to queens across Polistine wasps^69^. Comparisons between worker and gyne *P. metricus* found lower levels of *apolipophorin III* in worker-compared to gyne-destined larvae^70^. In *P. canadensis*, workers have increased apolipophorin compared to queens^71^. The gene is also upregulated during usurpation attempts in the socially parasitic *P. sulcifer*, suggesting that gene may have links to aggression in *Polistes^72^*. Apidaecin is an antimicrobial peptide involved in immunity^73^ and shows markedly increased expression in losers following social interactions with a later decreases (Fig 5b), suggesting possible immune activation in response to receiving aggression.

*Vitellogenin (vg)* is classically recognized for its role as an egg-yolk protein, which has a conserved role in oogenesis across insects^74^. In paper wasp*s*, levels of *vg* in the head or brain have been associated with social status, being highest in single and dominant foundresses and lowest in subordinate foundresses and workers^39,69,71,75^. Our data suggest that *vg* levels quickly respond to social interaction, rising substantially in both losers and winners relative to nonsocial controls at the early time point (Fig 5b). Winners maintain high levels of *vg* for hours after the interactions, while levels plummet in losers below those seen in nonsocial controls. By contrast, winners maintain high levels of *vg* following social interactions. Nonsocial control wasps show relatively lower levels of *vg* compared to socially interacting wasps, though it is hard to contextualize the *vg* levels observed in control wasps compared to those reported in other studies. Previous studies have examined patterns of gene expression in wasps in relation to life history state or broader social contexts (e.g. foundresses versus worker) and not in response to specific social experiences^39,69,71,75^. Additionally, the wasps in this study had been kept in the lab without nests following other studies of staged aggression contests^36,37,44^, which likely influences baseline levels of gene expression. Nevertheless, we find that *vg* is strongly upregulated in response to social interactions in general, but expression levels then diverge depending on social outcomes. To the extent that *vitellogenin* influences levels of aggression, the decrease seen over time in losers in this study may be indicative of a shift toward a submissive behavioral state.

We observed multiple genes that show increases in expression over time in losers in the central brain. The most upregulated gene in terms of log2 fold change in losers is a *myosin heavy chain* gene, which are upregulated in social wasp worker brains compared to queens^69^. We also observed a pattern of upregulation of *arrestin* in late losers but down regulation in winners and control nonsocial wasps. Previous studies of caste differential expression in *P. canadensis* found that *arrestin* was upregulated in workers relative to queens ^71^, and it is found upregulated among foragers in ants as well^76^. *FPPS* encodes farnesyl pyrophosphate synthase which is involved in JH production^77,78^ and is upregulated in queens in Polistine wasps^69^. We also observed increases in *inositol monophosphatase (imp)*, which is involved in the inositol phosphate signaling pathway ^79^ and has been linked to task differentiation in ants and bees^80,81^. Losers in our experiment would potentially become subordinate foundresses in a natural nesting context and not workers, though subordinates do more foraging than dominants^32^. Despite reduced reproduction and greater foraging relative to dominant foundresses, subordinate foundresses are not the same as workers and have been shown to have distinct neurogenomic profiles compared to dominant foundresses and workers in microarray and candidate-gene studies^39,66,75,82^. Nevertheless, the expression patterns of these genes suggest that within a few hours of emerging from a social encounter as a subordinate, multiple genes are dynamically regulated in a manner suggesting changes to aggression, reproduction, and metabolism (Fig 5).

Winners showed less extreme changes in gene expression over time compared to losers in our dataset (Fig 5). Among the genes with largest change by magnitude in winners are two members of *takeout* gene family, which show substantial decreases in losers (Fig 5). The *takeout* gene family is found across insects ^83^ and are they frequently regulated by juvenile hormone^84–86^. Both winners and losers showed increases in *Nieman Pick Type C2 (NPC2)*, which regulate steroid hormone biosynthesis including juvenile hormone^87^, and has been implicated in social communication among ant workers^88^. Notably, all three of these genes are among the most highly and significantly upregulated genes in the brain in response to social interactions (Fig 2a, 5b). The significant upregulation of these genes in response to social interactions and divergent patterns of expression between winners and losers over time make them interesting candidates for further study.

## CONCLUSIONS

The analysis of 139 RNAseq samples from the optic lobes and central brains of *P. fuscatus* foundresses revealed novel insights into the dynamic changes in neurogenomic states in peripheral and central nervous tissues following social interactions. Female *P. fuscatus* paper wasps have variable facial patterns that they use to visually recognize each other as individuals^5,37^. Though we did detect some differentially expressed genes in the optic lobe transcriptome in response to social interactions, changes in the brain were much larger and more dynamic, likely reflecting the importance of processing socially relevant information in more central brain regions as a key factor in driving neurogenomic shifts. After a 45-minute interaction, winners and losers show similar average changes in patterns of gene expression relative to nonsocial individuals, which may reflect the fact that the same neural circuits likely process initial social interactions regardless of the outcome. This result mirrors recent findings of similar neural firing patterns during social interactions in rodents and bats^21,22^ and similar neurogenomic responses shortly after fights in beta fish^28^.

Over a span of 4 hours the initial similarity between winners and losers decreases, as loser gene expression patterns show larger shifts consistent with theoretical predictions of larger loser effects compared to winner effects^68^. The most dramatic shifts in expression over the course of four hours in losers are due to a mixture of increasing or decreasing expression compared over time (Fig 5b). These data suggest that within a few hours a single subordinate experience can influence expression of multiple genes associated with behavioral and physiological differences, perhaps most notably *vitellogenin*. We do not suggest that a single social experience is necessarily sufficient to make a wasp into a subordinate foundress. Paper wasps engage in aggressive interactions on and off the nest early in the nesting cycle^32^ and many wasps that go on to become solitary or dominant foundresses likely experience some social defeats during this phase. Repeated interactions between co-nesting foundresses, however, are to likely compound and reinforce the types of effects we observe. Neurogenomic studies show shifts in neurogenomic profiles in many caste-associated genes in response to repeated wins or losses in dominance contests in *Ceratina* carpenter bees^11,24^. Paper wasps are notably plastic, with aggressive and dominant workers becoming more queenlike in the span of a few days when reproductive opportunities become available through experimental removal of the queen^26,27^. Moving forward, a major challenge is to understand how social experiences are processed in the brain giving rise to neurogenomic shifts and changes in expression of key regulators of behavior such as *vitellogenin*. Specifically, single-cell RNAseq approaches have the potential to indicate which cell-types are most strongly influenced by social interaction and could reveal how diverging gene expression patterns give rise to broader physiological consequences associated with social status.

## METHODS

### Experimental design and behavioral scoring

We tested the role of social experience on neurogenomic states comparing the responses of individuals to staged contests in a neutral arena to solitary experiences in the same arenas. Subjects were 90 female *P. fuscatus* collected during the pre-worker colony phase from their nests or while foraging in Tompkins county, New York in the spring of 2018 (Table S1). Wasps were brought into the lab and provided housing in small deli cups with *ad libitum* access to sugar and water. Prior to the trials, wasps were given identifying paint marks using Tester’s enamel paint to facilitate scoring of social interactions. During the trials, wasps were placed in a small neutral arena (100 mm diameter clear petri dish) with a plexiglass-lid under bright full spectrum lights either alone or with another wasp. Social trials featured pairings between weight-matched wasps that had been collected at distinct locations at least 2 kilometers apart, which is greater than the typical dispersal distances for this population^89^. While in the arenas, wasps were filmed for 45 minutes and then removed from the arenas. In half of the trials, wasps were immediately sacrificed by decapitation and their heads were placed in RNAlater for subsequent analysis. To aid uptake of RNAlater, small cuts were made on the exoskeleton of the head avoiding damaging neural tissue. In the other half of the trials, the wasps were returned to their individual housing and sacrificed 4 hours later using the same protocol. This generated four sets of samples: early social wasps (n = 30 wasps from 15 trials), early nonsocial wasps (n = 15), late social wasps (n = 30), and late nonsocial wasps (n=15, Fig 1a).

Videos of the social wasps were scored for stereotyped paper wasp aggressive behaviors including mounting, biting, hitting, grappling and darting^32,41^. Additionally, we scored when one wasp chased the other as an aggressive act. On average there were 33.13 ± 12.58 aggressive acts per trial. We categorized outcomes of encounters as either a win or loss based on the relative level of aggressive acts and whether or not one wasp mounted the other, a ritualized dominance behavior^32^.

### RNA sequencing and read processing

Brains were dissected from RNAlater-preserved wasp heads under a stereomicroscope. Optic lobes were separated from the rest of the brain (Fig 1b) and then combined for processing. We refer to these two tissue segments simply as the optic lobe and brain respectively in the text. RNA was extracted separately from the brain and combined optic lobes generating two pools of RNA from each wasp. Extracted RNA samples were sent to the Cornell Genomics Core for 3’RNA library preparation using the Lexogen kit. Due to low and/or poor-quality RNA yields for some samples, we were able to sequence 168 samples out of the intended 180. We sequenced libraries to an average coverage of 5.17 million single end 50 bp reads on a NextSeq500. Samples with less than 1 million reads were excluded from analyses due to their relatively low coverage, resulting in a final group of 139 RNAseq samples for analysis (Table S1).

We mapped reads to the *P. fuscatus* genome^42^ using STAR^90^. Read counts were calculated using HTseq with default settings^91^. Initial read counts revealed that the annotation of the *P. fuscatus* genome did not capture many 3’ untranslated regions, so we manually scanned the genome to update gene body annotations. To identify 3’ untranslated regions we jointly visualized paired-end mRNAseq reads from female *P. fuscatus* heads with a sample of 3’ RNAseq reads using the Integrated Genome Viewer^92^ and updated a GTF file based on this scan. In addition to extending the UTRs, in some cases we combined genes, separated genes or identified genes not previously included in the prior annotation. The GTF file used for this study is provided as a supplemental file. Before engaging in downstream differential expression analyses, we first inspected the separation of the samples using principal component analysis (PCA) to ensure that brain and optic lobe tissues had distinct expression profiles, as would be expected based on differential cellular composition of the samples. The PCA was calculated by using the ‘vst’ normalization function of DESeq2^93^. Inspection of the samples plotted against PC1 and PC2 revealed 2 distinct clusters of samples corresponding to optic lobe and brain respectively (Fig 1b). Additionally, we removed non-expressed or lowly-expressed genes from the count table in order to make analyses faster. After filtering, we were left with 8219 genes for further analyses.

### Gene expression analyses

Patterns of differential expression were determined using DESeq2^93^ in R v 3.6.2 (R Team 2019). Depending on the analysis we examined the entire data set (both brain and optic lobes), only the brain data or only the optic lobe data using linear models with fixed effects. All R code used for analysis is provided. First, we considered models with social experience treatment (social v. nonsocial), tissue (brain v. optic lobe) and time (early v. late). We examined the interactive effects following the recommendations of the authors of the DESeq2 analysis package ^93^. We generated combined variables to examine differences in expression across groups. For example, to look at the effects of time and social experience we classified samples as belonging to one of four groups early_social, early_nonsocial, late_social, or late_nonsocial under a single categorical variable, e.g., time_social. By comparing contrasts among the different pairs of categories, we were able to determine how different combinations of samples influence patterns of differential gene expression. For analyses looking at contest outcome, we only examined social trials for which at least 10 aggressive acts occurred. The outcome of the trial was coded as winner, loser or nonsocial. Genes were considered to be differentially expressed if the FDR adjusted P value ≤ 0.1.

We compared patterns of differential expression in winners and losers, based on log2 fold changes in expression. Since absolutely small changes in lowly expressed genes can give rise to large log2 fold changes, we first removed all genes with mean expression below 100 before comparing patterns of expression. First, we compared expression relative to nonsocial wasps in winners and losers respectively in the combined brain and optic lobe datasets. In analyses focusing solely on the brain dataset, we examined how the relations between winner and loser expression profiles changed between early and late sampling points using a general linear mixed model implemented in the lme4 package for R^95^. Log2 fold change differences in expression relative to nonsocial wasps sacrificed at the same time were used as a basis of comparison. We modeled relative fold change in losers as a function of the relative fold change in winners, time, their interaction, and gene ID as a random effect. We also separately examined the relationship between winners and losers at early and late time points using a linear model. Finally, we compared the relative log2 fold changes between the earlier and later time points for losers to the changes observed in winners. In these comparisons positive values of expression denote increased expression at the later time point.

Genes are frequently expressed in a modular manner, with groups of genes showing similar expression patterns^63^, so we calculated co-expression modules from our brain dataset using WGCNA. This analysis focused on understanding modules associated with winning or losing at different time points, so we limited our analysis to a subset of the brain RNAseq data set that had engaged in more vigorous encounters (i.e., winner, loser and nonsocial). R code used for analysis is provided as a supplemental file.

### Gene ontology

We used gene ontology enrichment analyses to identify gene functions that were enriched in our various parts of our dataset. The *P. fuscatus* gene set was annotated using the Blast2GO function of OmicsBox based on sequence similarity with *Drosophila melanogaster* genes^96^. For enrichment analyses, we used the TopGO package in R^97^. We only included categories with at least 10 annotated genes in the dataset. Significantly over-represented categories were identified using the ‘weigh01’ function in TopGO with the ‘classicfisher’ statistic.

## Data Availability

Raw sequence data are available in the NCBI Short Read Archive under PRJNA705303.

## Funding

This research was funded by grant NIH-DP2-GM128202 to MJS.

## Author contributions

FMK, CMJ and MJS designed the study. FMK and CMJ conducted behavioral trials. FMK processed tissue samples and extracted RNA. NZ scored behavioral trials. SEM, NZ, EM and MJS processed and aligned RNA data. MJS analyzed the data and wrote the manuscript with input from the other authors.

## Notes

### Competing Interest Statement

The authors have declared no competing interest.

## REFERNCES

1. Ligon, R. A. Defeated chameleons darken dynamically during dyadic disputes to decrease danger from dominants. Behav. Ecol. Sociobiol. 68, 1007–1017 (2014).

2. Chandrasekaran, S. et al. Aggression is associated with aerobic glycolysis in the honey bee brain1. Genes Brain Behav. 14, 158–166 (2015).

3. Rittschof, C. C. & Hughes, K. A. Advancing behavioural genomics by considering timescale. Nat. Com mu n. 9, 1–11 (2018).

4. Snyder-Mackler, N. et al. Social determinants of health and survival in humans and other animals. Science 368, (2020).

5. Sheehan, M. J. & Tibbetts, E. A. Specialized face learning is associated with individual recognition in paper wasps, science 334, 1272–1275 (2011).

6. Rittschof, C. C. et al. Neuromolecular responses to social challenge: Common mechanisms across mouse, stickleback fish, and honey bee. Proc. Natl. Acad. Sci. 111, 17929–17934 (2014).

7. Dugatkin, L. A. & Earley, R. L. Group fusion: the impact of winner, loser, and bystander effects on hierarchy formation in large groups. Behav. Ecol. 14, 367–373 (2003).

8. Hsu, Y. Y., Lee, I. H. & Lu, C. K. Prior contest information: mechanisms underlying winner and loser effects. Behav. Ecol. Sociobiol. 63, 1247–1257 (2009).

9. Lehner, S. R., Rutte, C. & Taborsky, M. Rats benefit from winner and loser effects. Ethology 117, 949–960 (2011).

10. Trannoy, S., Penn, J., Lucey, K., Popovic, D. & Kravitz, E. A. Short and long-lasting behavioral consequences of agonistic encounters between male Drosophila melanogaster. Proc. Natl. Acad. Sci. 113, 4818–4823 (2016).

11. Withee, J. R. & Rehan, S. M. Social aggression, experience, and brain gene expression in a subsocial bee. Integr. Comp. Biol. 57, 640–648 (2017).

12. Harrison, L. M., Jennions, M. D. & Head, M. L. Does the winner-loser effect determine male mating success? Biol. Lett. 14, 20180195 (2018).

13. Tibbetts, E. A., Fearon, M. L., Wong, E., Huang, Z. Y. & Tinghitella, R. M. Rapid juvenile hormone downregulation in subordinate wasp queens facilitates stable cooperation. Proc. R. Soc. B Biol. Sci. 285, 20172645 (2018).

14. Reeve, H. K., Starks, P. T., Peters, J. M. & Nonacs, P. Genetic support for the evolutionary theory of reproductive transactions in social wasps. Proc. R. Soc. Lond. B Biol. Sci. 267, 75–79 (2000).

15. Seppa, P., Queller, D. C. & Strassmann, J. E. Reproduction in foundress associations of the social wasp, Polistes Carolina: conventions, competition, and skew. Behav. Ecol. 13, 531–542 (2002).

16. Gospocic, J. et al. The neuropeptide corazonin controls social behavior and caste identity in ants. Cell 170, 748–759. e12 (2017).

17. Razzoli, M. et al. Social stress shortens lifespan in mice. Aging Cell 17, (2018).

18. Bukhari, S. A. et al. Temporal dynamics of neurogenomic plasticity in response to social interactions in male threespined sticklebacks. PLoS Genet. 13, e1006840 (2017).

19. Saul, M. C. et al. Transcriptional regulatory dynamics drive coordinated metabolic and neural response to social challenge in mice. Genome Res. 27, 959–972 (2017).

20. O’Connell, L. A. & Hofmann, H. A. Evolution of a vertebrate social decision-making network. Science 336, 1154–1157 (2012).

21. Kingsbury, L. et al. Correlated neural activity and encoding of behavior across brains of socially interacting animals. Cell 178, 429–446. e16 (2019).

22. Zhang, W. & Yartsev, M. M. Correlated neural activity across the brains of socially interacting bats. Cell 178, 413–428. e22 (2019).

23. Oliveira, R. F. et al. Assessment of fight outcome is needed to activate socially driven transcriptional changes in the zebrafish brain. Proc. Natl. Acad. Sci. 113, E654–E661 (2016).

24. Steffen, M. A. & Rehan, S. M. Genetic signatures of dominance hierarchies reveal conserved cis-regulatory and brain gene expression underlying aggression in a facultatively social bee. Genes Brain Behav. 19, e12597 (2020).

25. Opachaloemphan, C. et al. Early behavioral and molecular events leading to caste switching in the ant Harpegnathos. Genes Dev. (2021).

26. Taylor, B. A., Cini, A., Cervo, R., Reuter, M. & Sumner, S. Queen succession conflict in the paper wasp Polistes dominula is mitigated by age-based convention. Behav. Ecol. 31, 992–1002 (2020).

27. Taylor, B. A., Cini, A., Wyatt, C. D., Reuter, M. & Sumner, S. The molecular basis of socially mediated phenotypic plasticity in a eusocial paper wasp. Nat. Commun. 12, 1–10 (2021).

28. Vu, T.-D. et al. Behavioral and brain-transcriptomic synchronization between the two opponents of a fighting pair of the fish Betta splendens. PLoS Genet. 16, e1008831 (2020).

29. Reeve, H. K. Polistes. in The Social Biology of Wasps 99–148 (Comstock, 1991).

30. Jandt, J. M., Tibbetts, E. A. & Toth, A. L. Polistes paper wasps: a model genus for the study of social dominance hierarchies. Insectes Sociaux 61, 11–27 (2014).

31. Sheehan, M. J. et al. Different axes of environmental variation explain the presence vs. extent of cooperative nest founding associations in Polistes paper wasps. Ecol. Lett. 18, 1057–1067 (2015).

32. West Eberhard, M. J. The Social Biology of Polistine Wasps. (Museum of Zoology, University of Michigan, 1969).

33. Gamboa, G. J., Wacker, T. L., Duffy, K. G., Dobson, S. W. & Fishwild, T. G. Defense against intraspecific usurpation by paper wasp cofoundresses (Polistes fuscatus, Hymenoptera, Vespidae). Can. J. Zool.-Rev. Can. Zool. 70, 2369–2372 (1992).

34. Tibbetts, E. A. & Reeve, H. K. Aggression and resource sharing among foundresses in the social wasp Polistes dominulus: testing transactional theories of conflict. Behav. Ecol. Sociobiol. 48, 344–352 (2000).

35. Mora-Kepfer, F. Context-dependent acceptance of non-nestmates in a primitively eusocial insect. Behav. Ecol. Sociobiol. 68, 363–371 (2014).

36. Tibbetts, E. A. & Dale, J. A socially enforced signal of quality in a paper wasp. Nature 432, 218–222 (2004).

37. Sheehan, M. J. & Tibbetts, E. A. Evolution of identity signals: Frequency-dependent benefits of distinctive phenotypes used for individual recognition. Evolution 63, 3106–3113 (2009).

38. Berens, A. J., Tibbetts, E. A. & Toth, A. L. Candidate genes for individual recognition in Polistes fuscatus paper wasps. J. Comp. Physiol. A 202, 115–129 (2016).

39. Toth, A. L. et al. Shared genes related to aggression, rather than chemical communication, are associated with reproductive dominance in paper wasps (Polistes metricus). BMC Genomics 15, 75 (2014).

40. Tibbetts, E. A. Visual signals of individual identity in the wasp Polistes fuscatus. Proc. R. Soc. Lond. B Biol. Sci. 269, 1423–1428 (2002).

41. Sheehan, M. J. & Tibbetts, E. A. Selection for individual recognition and the evolution of polymorphic identity signals in Polistes paper wasps. J. Evol. Biol. 23, 570–577 (2010).

42. Miller, S. E. et al. Evolutionary dynamics of recent selection on cognitive abilities. Proc. Natl. Acad. Sci. 117, 3045–3052 (2020).

43. Berens, A. J., Tibbetts, E. A. & Toth, A. L. Cognitive specialization for learning faces is associated with shifts in the brain transcriptome of a social wasp. J. Exp. Biol. 220, 2149–2153 (2017).

44. Sheehan, M. J. & Tibbetts, E. A. Robust long-term social memories in a paper wasp. Curr. Biol. CB 18, R851–R852 (2008).

45. Li, L. et al. Large-scale transcriptome changes in the process of long-term visual memory formation in the bumblebee, Bombus terrestris. Sci. Rep. 8, 1–10 (2018).

46. Tibbetts, E. A., Wong, E. & Bonello, S. Wasps use social eavesdropping to learn about individual rivals. Curr. Biol. 30, 3007–3010. e2 (2020).

47. Sisneros, J. A. & Bass, A. H. Seasonal plasticity of peripheral auditory frequency sensitivity. J. Neurosci. 23, 1049–1058 (2003).

48. Dey, S. et al. Cyclic regulation of sensory perception by a female hormone alters behavior. Cell 161, 1334–1344 (2015).

49. Walker, S. J., Corrales-Carvajal, V. M. & Ribeiro, C. Postmating circuitry modulates salt taste processing to increase reproductive output in Drosophila. Curr. Biol. 25, 2621–2630 (2015).

50. van der Linden, C., Jakob, S., Gupta, P., Dulac, C. & Santoro, S. W. Sex separation induces differences in the olfactory sensory receptor repertoires of male and female mice. Nat. Commun. 9, 1–15 (2018).

51. Butler, J. M. et al. Reproductive state-dependent plasticity in the visual system of an African cichlid fish. Horm. Behav. 114, 104539 (2019).

52. Veen, T., Brock, C., Rennison, D. & Bolnick, D. Plasticity contributes to a fine-scale depth gradient in sticklebacks’ visual system. Mol. Ecol. 26, 4339–4350 (2017).

53. Tibbetts, E. A., Desjardins, E., Kou, N. & Wellman, L. Social isolation prevents the development of individual face recognition in paper wasps. Anim. Behav. 152, 71–77 (2019).

54. Alekseyenko, O. V. et al. Serotonergic modulation of aggression in Drosophila involves GABAergic and cholinergic opposing pathways. Curr. Biol. 29, 2145–2156. e5 (2019).

55. Palavicino-Maggio, C. B., Chan, Y.-B., McKellar, C. & Kravitz, E. A. A small number of cholinergic neurons mediate hyperaggression in female Drosophila. Proc. Natl. Acad. Sci. 116, 17029–17038 (2019).

56. Schretter, C. E. et al. Cell types and neuronal circuitry underlying female aggression in Drosophila. Elife 9, e58942 (2020).

57. Tully, T., Preat, T., Boynton, S. C. & Del Vecchio, M. Genetic dissection of consolidated memory in Drosophila. Cell 79, 35–47 (1994).

58. McGaugh, J. L. Memory-a century of consolidation. Science 287, 248–251 (2000).

59. Mery, F. & Kawecki, T. J. A cost of long-term memory in Drosophila. Science 308, 1148–1148 (2005).

60. Moncada, D. & Viola, H. Induction of long-term memory by exposure to novelty requires protein synthesis: evidence for a behavioral tagging. J. Neurosci. 27, 7476–7481 (2007).

61. Jarome, T. J. & Helmstetter, F. J. Protein degradation and protein synthesis in long-term memory formation. Front. Mol. Neurosci. 7, 61 (2014).

62. Kandel, E. R., Dudai, Y. & Mayford, M. R. The molecular and systems biology of memory. Cell 157, 163–186 (2014).

63. Langfelder, P. & Horvath, S. WGCNA: an R package for weighted correlation network analysis. BMC Bioinformatics 9, 559 (2008).

64. Creel, S. Dominance, aggression, and glucocorticoid levels in social carnivores. J. Mammal. 86, 255–264 (2005).

65. Williamson, C. M., Lee, W., Romeo, R. D. & Curley, J. P. Social context-dependent relationships between mouse dominance rank and plasma hormone levels. Physiol. Behav. 171, 110–119 (2017).

66. Toth, A. L. et al. Brain transcriptomic analysis in paper wasps identifies genes associated with behaviour across social insect lineages. Proc. R. Soc. Lond. B Biol. Sci. 277, 2139–2148 (2010).

67. Cardoso, S. D., Teles, M. C. & Oliveira, R. F. Neurogenomic mechanisms of social plasticity. J. Exp. Biol. 218, 140–149 (2015).

68. Leimar, O. The evolution of social dominance through reinforcement learning. bioRxiv (2020).

69. Wyatt, C. D. et al. Genetic toolkit for sociality predicts castes across the spectrum of social complexity in wasps. bioRxiv (2020).

70. Hunt, J. H. et al. Differential gene expression and protein abundance evince ontogenetic bias toward castes in a primitively eusocial wasp. PloS One 5, e10674 (2010).

71. Sumner, S., Pereboom, J. J. & Jordan, W. C. Differential gene expression and phenotypic plasticity in behavioural castes of the primitively eusocial wasp, Polistes canadensis. Proc. R. Soc. Lond. B Biol. Sci. 273, 19–26 (2006).

72. Cini, A. et al. Social parasitism and the molecular basis of phenotypic evolution. Front. Genet. 6, 32 (2015).

73. Casteels, P., Ampe, C., Jacobs, F., Vaeck, M. & Tempst, P. Apidaecins: antibacterial peptides from honeybees. EMBOJ. 8, 2387–2391 (1989).

74. Hagedorn, H. H. & Kunkel, J. G. Vitellogenin and vitellin in insects. Annu. Rev. Entomol. 24, 475–505 (1979).

75. Manfredini, F., Brown, M. J. & Toth, A. L. Candidate genes for cooperation and aggression in the social wasp Polistes dominula. J. Comp. Physiol. A 204, 449–463 (2018).

76. Feldmeyer, B., Elsner, D. & Foitzik, S. Gene expression patterns associated with caste and reproductive status in ants: worker-specific genes are more derived than queen-specific ones. Mol. Ecol. 23, 151–161 (2014).

77. Tsang, S. S. et al. Diversity of insect sesquiterpenoid regulation. Front. Genet. 11, (2020).

78. Bomtorin, A. D. et al. Juvenile hormone biosynthesis gene expression in the corpora allata of honey bee (Apis mellifera L.) female castes. PloS One 9, e86923 (2014).

79. Berridge, M. J. Inositol trisphosphate and calcium signalling mechanisms. Biochim. Biophys. Acta BBA-Mol. Cell Res. 1793, 933–940 (2009).

80. Lutz, C. C., Rodriguez-Zas, S. L., Fahrbach, S. E. & Robinson, G. E. Transcriptional response to foraging experience in the honey bee mushroom bodies. Dev. Neurobiol. 72, 153–166 (2012).

81. Friedman, D. A. et al. The role of dopamine in the collective regulation of foraging in harvester ants. Iscience 8, 283–294 (2018).

82. Toth, A. L. et al. Wasp gene expression supports an evolutionary link between maternal behavior and eusociality. Science 318, 441–444 (2007).

83. Vanaphan, N., Dauwalder, B. & Zufall, R. A. Diversification of takeout, a male-biased gene family in Drosophila. Gene 491, 142–148 (2012).

84. Du, J., Hiruma, K. & Riddiford, L. M. A novel gene in the takeout gene family is regulated by hormones and nutrients in Manduca larval epidermis. Insect Biochem. Mol. Biol. 33, 803–814 (2003).

85. Hagai, T., Cohen, M. & Bloch, G. Genes encoding putative Takeout/juvenile hormone binding proteins in the honeybee (Apis mellifera) and modulation by age and juvenile hormone of the takeout-like gene GB19811. Insect Biochem. Mol. Biol. 37, 689–701 (2007).

86. Chamseddin, K. H. et al. takeout-dependent longevity is associated with altered Juvenile Hormone signaling. Mech. Ageing Dev. 133, 637–646 (2012).

87. Huang, X., Warren, J. T., Buchanan, J., Gilbert, L. I. & Scott, M. P. Drosophila Niemann-Pick type C-2 genes control sterol homeostasis and steroid biosynthesis: a model of human neurodegenerative disease. Development 134, 3733–3742 (2007).

88. Ishida, Y. et al. Niemann-Pick type C2 protein mediating chemical communication in the worker ant. Proc. Natl. Acad. Sci. 111, 3847–3852 (2014).

89. Bluher, S. E., Miller, S. E. & Sheehan, M. J. Fine-scale population structure but limited genetic differentiation in a cooperatively breeding paper wasp. Genome Biol. Evol. 12, 701–714 (2020).

90. Dobin, A. et al. STAR: ultrafast universal RNA-seq aligner. Bioinformatics 29, 15–21 (2013).

91. Anders, S., Pyl, P. T. & Huber, W. HTSeq—a Python framework to work with high-throughput sequencing data. Bioinformatics 31, 166–169 (2015).

92. Robinson, J. T., Thorvaldsdóttir, H., Wenger, A. M., Zehir, A. & Mesirov, J. P. Variant review with the integrative genomics viewer. Cancer Res. 77, e31–e34 (2017).

93. Love, M. I., Huber, W. & Anders, S. Moderated estimation of fold change and dispersion for RNA-seq data with DESeq2. Genome Biol. 15, 550 (2014).

94. Team, R. C. R:A language and environment for statistical computing. (Vienna, Austria, 2013).

95. Bates, D., Sarkar, D., Bates, M. D. & Matrix, L. The lme4 package. R Package Version 2, 74 (2007).

96. Conesa, A. et al. Blast2GO: a universal tool for annotation, visualization and analysis in functional genomics research. Bioinformatics 21, 3674–3676 (2005).

97. Alexa, A. & Rahnenführer, J. Gene set enrichment analysis with topGO. Bioconductor Improv 27, (2009).

