## Supplemental files for "Dynamic neurogenomic responses to social interactions and dominance outcomes in female paper wasps"

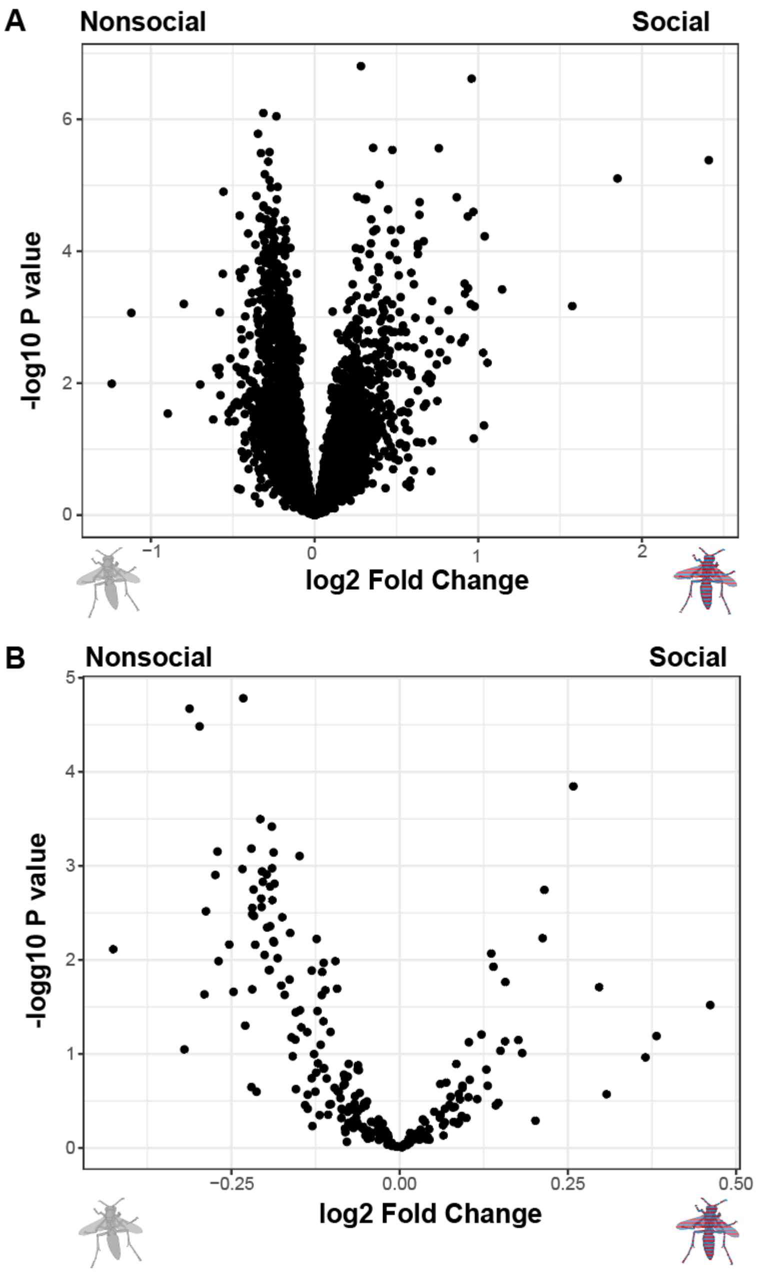

**Figure S1: Volcano plot of socially regulated genes in full dataset**

The volcano plots show the genes that are differentially expressed between individuals that experiences social interaction (social) versus those that did not (nonsocial). Higher log2 fold change values indicate upregulation in the social group compared to the nonsocial group. Panel A shows the data for all genes examined. Panel B shows the plot for genes annotated with memory-related functions. The red and blue striped wasp symbol indicates that the data includes all socially interacting wasps

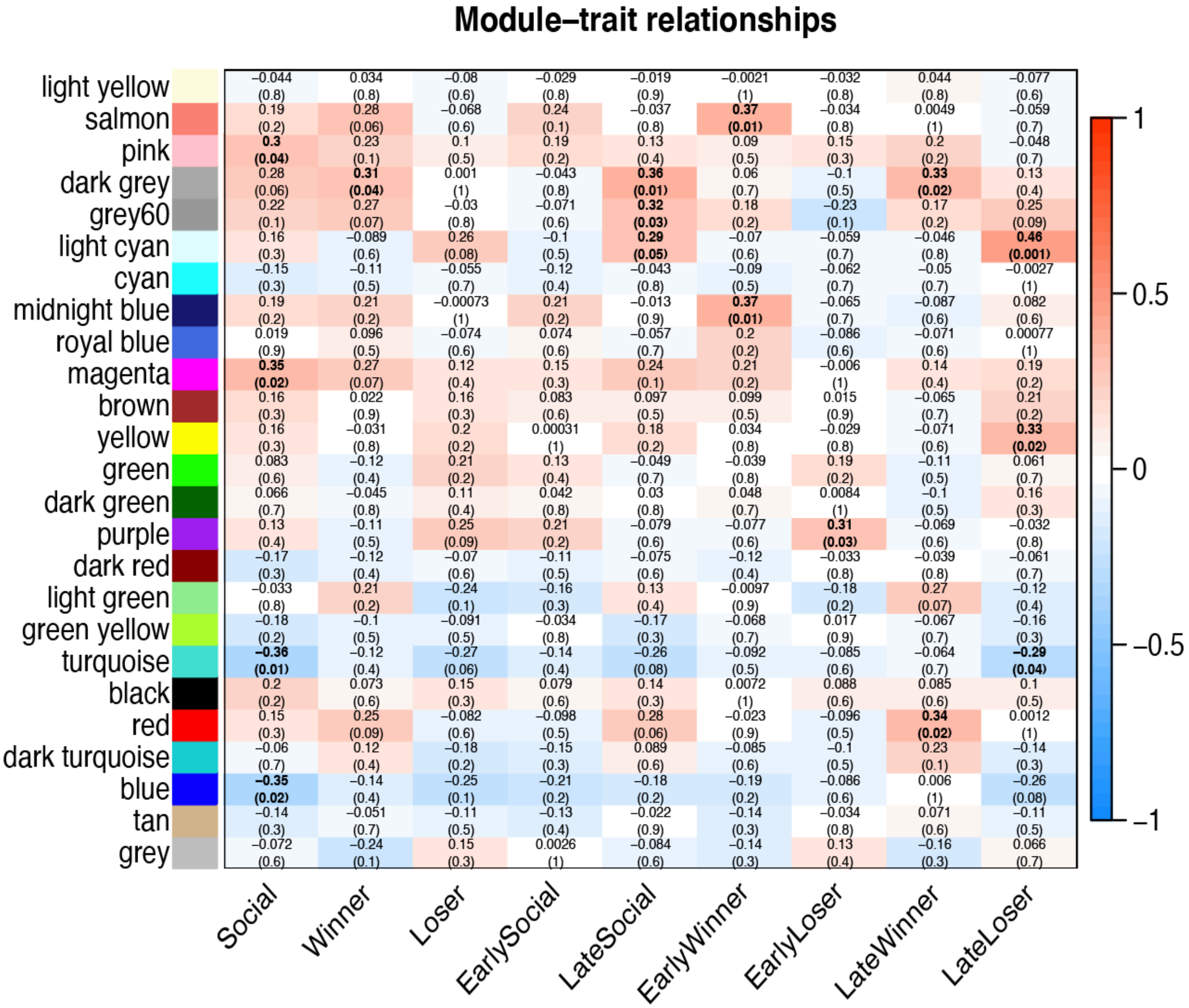

**Figure S2: Module outcome relationships in the brain dataset**

The heatmap shows the relationships between the modules of co-expressed genes identified by WGCNA with different social outcomes. White and lighter colors indicate low correlations. Bolder reds indicate stronger positive correlations. Bolder blues indicate stronger negative correlations. In each cell the top line of text reports the correlation between a given module and the social trait or time since social interaction. The bottom line of text shows the P value in parentheses. Significant correlations ($P \leq 0.05$) are highlighted with bold text.

**Table S1**

| Sample | Time | Social | Tissue | Outcome | Sequence Depth |
|---|---|---|---|---|---|
| Fuscatus_2018_101A | Early | Nonsocial | Optic | Nonsocial | 5280578 |
| Fuscatus_2018_101B | Early | Nonsocial | Brain | Nonsocial | 2738780 |
| Fuscatus_2018_104A | Late | Nonsocial | Optic | Nonsocial | 7103006 |
| Fuscatus_2018_104B | Late | Nonsocial | Brain | Nonsocial | 3606890 |
| Fuscatus_2018_106A | Early | Nonsocial | Optic | Nonsocial | 2364946 |
| Fuscatus_2018_106B | Early | Nonsocial | Brain | Nonsocial | 3540467 |
| Fuscatus_2018_121A | Early | Social | Optic | Loser | 4344918 |
| Fuscatus_2018_121B | Early | Social | Brain | Loser | 4552113 |
| Fuscatus_2018_126A | Early | Social | Optic | Winner | 4574347 |
| Fuscatus_2018_126B | Early | Social | Brain | Winner | 5603436 |
| Fuscatus_2018_142A | Late | Social | Optic | Loser | 4311099 |
| Fuscatus_2018_14A | Late | Social | Optic | Loser | 5278364 |
| Fuscatus_2018_14B | Late | Social | Brain | Loser | 5839144 |
| Fuscatus_2018_151A | Late | Nonsocial | Optic | Nonsocial | 5865655 |
| Fuscatus_2018_151B | Late | Nonsocial | Brain | Nonsocial | 4432685 |
| Fuscatus_2018_152A | Late | Nonsocial | Optic | Nonsocial | 4191333 |
| Fuscatus_2018_152B | Late | Nonsocial | Brain | Nonsocial | 2897874 |
| Fuscatus_2018_156A | Early | Social | Brain | Loser | 6378977 |
| Fuscatus_2018_156B | Early | Social | Optic | Loser | 7508942 |
| Fuscatus_2018_165A | Early | Social | Optic | Winner | 6615619 |
| Fuscatus_2018_167A | Late | Social | Optic | Loser | 6504306 |
| Fuscatus_2018_167B | Late | Social | Brain | Loser | 5936427 |
| Fuscatus_2018_181A | Early | Social | Optic | Loser | 4527025 |
| Fuscatus_2018_181B | Early | Social | Brain | Loser | 4577560 |
| Fuscatus_2018_183A | Late | Social | Optic | Winner | 8798872 |
| Fuscatus_2018_183B | Late | Social | Brain | Winner | 4454285 |
| Fuscatus_2018_185A | Late | Social | Optic | Winner | 4997432 |
| Fuscatus_2018_185B | Late | Social | Brain | Winner | 2275757 |
| Fuscatus_2018_186A | Late | Social | Optic | NA_low aggression in trial | 6410919 |
| Fuscatus_2018_186B | Late | Social | Brain | NA_low aggression in trial | 5899077 |
| Fuscatus_2018_18A | Early | Social | Optic | NA_low aggression in trial | 3928437 |
| Fuscatus_2018_18B | Early | Social | Brain | NA_low aggression in trial | 5298738 |
| Fuscatus_2018_273A | Late | Nonsocial | Optic | Nonsocial | 5973631 |
| Fuscatus_2018_273B | Late | Nonsocial | Brain | Nonsocial | 2054190 |
| Fuscatus_2018_274A | Late | Social | Optic | Winner | 4414126 |
| Fuscatus_2018_274B | Late | Social | Brain | Winner | 3920857 |
| Fuscatus_2018_275A | Early | Social | Optic | Winner | 7765170 |
| Fuscatus_2018_281A | Late | Social | Optic | Winner | 8377327 |

| | | | | | |
|---|---|---|---|---|---|
| Fuscatus_2018_281B | Late | Social | Brain | Winner | 6650264 |
| Fuscatus_2018_283A | Late | Social | Optic | NA_low aggression in trial | 5580202 |
| Fuscatus_2018_283B | Late | Social | Brain | NA_low aggression in trial | 5330802 |
| Fuscatus_2018_287A | Late | Social | Optic | NA_low aggression in trial | 4344088 |
| Fuscatus_2018_287B | Late | Social | Brain | NA_low aggression in trial | 12146881 |
| Fuscatus_2018_293A | Early | Social | Optic | Winner | 4664752 |
| Fuscatus_2018_293B | Early | Social | Brain | Winner | 4263418 |
| Fuscatus_2018_294A | Early | Social | Optic | Winner | 4034741 |
| Fuscatus_2018_298A | Early | Nonsocial | Optic | Nonsocial | 4584021 |
| Fuscatus_2018_298B | Early | Nonsocial | Brain | Nonsocial | 5080373 |
| Fuscatus_2018_318A | Late | Social | Optic | Loser | 5648198 |
| Fuscatus_2018_319A | Early | Social | Optic | Loser | 5048582 |
| Fuscatus_2018_319B | Early | Social | Brain | Loser | 4955401 |
| Fuscatus_2018_343A | Early | Social | Optic | Winner | 6255474 |
| Fuscatus_2018_343B | Early | Social | Brain | Winner | 5216375 |
| Fuscatus_2018_350A | Early | Nonsocial | Optic | Nonsocial | 5319770 |
| Fuscatus_2018_350B | Early | Nonsocial | Brain | Nonsocial | 3708204 |
| Fuscatus_2018_379A | Early | Social | Optic | Winner | 4136108 |
| Fuscatus_2018_379B | Early | Social | Brain | Winner | 4084916 |
| Fuscatus_2018_381A | Early | Social | Optic | Loser | 5442833 |
| Fuscatus_2018_382A | Late | Social | Optic | NA_low aggression in trial | 3584617 |
| Fuscatus_2018_382B | Late | Social | Brain | NA_low aggression in trial | 5831974 |
| Fuscatus_2018_38A | Early | Social | Optic | NA_low aggression in trial | 5498562 |
| Fuscatus_2018_38B | Early | Social | Brain | NA_low aggression in trial | 6712107 |
| Fuscatus_2018_39A | Early | Social | Optic | Winner | 8888570 |
| Fuscatus_2018_39B | Early | Social | Brain | Winner | 2622165 |
| Fuscatus_2018_400A | Late | Nonsocial | Optic | Nonsocial | 5731695 |
| Fuscatus_2018_400B | Late | Nonsocial | Brain | Nonsocial | 3547448 |
| Fuscatus_2018_40A | Late | Nonsocial | Optic | Nonsocial | 6370897 |
| Fuscatus_2018_40B | Late | Nonsocial | Brain | Nonsocial | 6628319 |
| Fuscatus_2018_415A | Early | Social | Optic | Loser | 6630046 |
| Fuscatus_2018_415B | Early | Social | Brain | Loser | 6952449 |
| Fuscatus_2018_41A | Early | Nonsocial | Optic | Nonsocial | 5474886 |
| Fuscatus_2018_41B | Early | Nonsocial | Brain | Nonsocial | 5736866 |
| Fuscatus_2018_429A | Late | Social | Optic | Winner | 4362571 |
| Fuscatus_2018_42A | Early | Social | Optic | NA_low aggression in trial | 3085552 |
| Fuscatus_2018_42B | Early | Social | Brain | NA_low aggression in trial | 4666995 |
| Fuscatus_2018_448B | Early | Social | Brain | Winner | 8114115 |
| Fuscatus_2018_451A | Early | Social | Optic | Loser | 6795091 |
| Fuscatus_2018_451B | Early | Social | Brain | Loser | 6390624 |

| | | | | | |
|---|---|---|---|---|---|
| Fuscatus_2018_453A | Late | Social | Optic | Winner | 5127231 |
| Fuscatus_2018_453B | Late | Social | Brain | Winner | 5076070 |
| Fuscatus_2018_457A | Late | Social | Optic | Winner | 6414076 |
| Fuscatus_2018_457B | Late | Social | Brain | Winner | 6269728 |
| Fuscatus_2018_460A | Late | Social | Optic | NA_low aggression in trial | 5064611 |
| Fuscatus_2018_460B | Late | Social | Brain | NA_low aggression in trial | 5002466 |
| Fuscatus_2018_481A | Early | Nonsocial | Brain | Nonsocial | 5238365 |
| Fuscatus_2018_481B | Early | Nonsocial | Optic | Nonsocial | 7427134 |
| Fuscatus_2018_484A | Late | Social | Brain | Loser | 2598848 |
| Fuscatus_2018_484B | Late | Social | Optic | Loser | 4156235 |
| Fuscatus_2018_485A | Late | Social | Optic | NA_low aggression in trial | 5245121 |
| Fuscatus_2018_487A | Early | Nonsocial | Optic | Nonsocial | 3397925 |
| Fuscatus_2018_487B | Early | Nonsocial | Brain | Nonsocial | 2994692 |
| Fuscatus_2018_488A | Early | Nonsocial | Optic | Nonsocial | 3492612 |
| Fuscatus_2018_490A | Early | Social | Optic | Loser | 5038856 |
| Fuscatus_2018_490B | Early | Social | Brain | Loser | 4530716 |
| Fuscatus_2018_493A | Late | Social | Optic | NA_low aggression in trial | 3497329 |
| Fuscatus_2018_493B | Late | Social | Brain | NA_low aggression in trial | 3643995 |
| Fuscatus_2018_494A | Late | Social | Optic | Loser | 4708046 |
| Fuscatus_2018_496A | Late | Nonsocial | Optic | Nonsocial | 8148044 |
| Fuscatus_2018_496B | Late | Nonsocial | Brain | Nonsocial | 6801794 |
| Fuscatus_2018_499A | Early | Nonsocial | Optic | Nonsocial | 4482612 |
| Fuscatus_2018_499B | Early | Nonsocial | Brain | Nonsocial | 5702849 |
| Fuscatus_2018_501B | Late | Social | Brain | Winner | 7417577 |
| Fuscatus_2018_515A | Early | Social | Brain | Loser | 2386525 |
| Fuscatus_2018_515B | Early | Social | Optic | Loser | 7530795 |
| Fuscatus_2018_530A | Early | Social | Optic | NA_low aggression in trial | 5743440 |
| Fuscatus_2018_530B | Early | Social | Brain | NA_low aggression in trial | 3089446 |
| Fuscatus_2018_531A | Early | Nonsocial | Optic | Nonsocial | 3768278 |
| Fuscatus_2018_531B | Early | Nonsocial | Brain | Nonsocial | 5944213 |
| Fuscatus_2018_532B | Early | Social | Brain | Loser | 3919356 |
| Fuscatus_2018_53A | Early | Social | Optic | Winner | 4680680 |
| Fuscatus_2018_53B | Early | Social | Brain | Winner | 5110080 |
| Fuscatus_2018_56A | Late | Social | Optic | Winner | 5818043 |
| Fuscatus_2018_56B | Late | Social | Brain | Winner | 7379130 |
| Fuscatus_2018_61B | Late | Nonsocial | Brain | Nonsocial | 5336389 |
| Fuscatus_2018_63A | Late | Nonsocial | Optic | Nonsocial | 5556741 |
| Fuscatus_2018_63B | Late | Nonsocial | Brain | Nonsocial | 5717866 |
| Fuscatus_2018_66A | Early | Nonsocial | Brain | Nonsocial | 4383935 |
| Fuscatus_2018_66B | Early | Nonsocial | Optic | Nonsocial | 5875286 |

| | | | | | |
|---|---|---|---|---|---|
| Fuscatus_2018_68A | Early | Social | Optic | NA_low aggression in trial | 2502864 |
| Fuscatus_2018_68B | Early | Social | Brain | NA_low aggression in trial | 3391640 |
| Fuscatus_2018_69A | Early | Social | Optic | NA_low aggression in trial | 5764656 |
| Fuscatus_2018_70A | Late | Nonsocial | Optic | Nonsocial | 2832267 |
| Fuscatus_2018_70B | Late | Nonsocial | Brain | Nonsocial | 3416810 |
| Fuscatus_2018_73A | Early | Social | Optic | NA_low aggression in trial | 3508618 |
| Fuscatus_2018_73B | Early | Social | Brain | NA_low aggression in trial | 6079034 |
| Fuscatus_2018_77B | Late | Social | Brain | Loser | 4568311 |
| Fuscatus_2018_80A | Late | Social | Optic | Loser | 5519797 |
| Fuscatus_2018_80B | Late | Social | Brain | Loser | 5696110 |
| Fuscatus_2018_81A | Early | Nonsocial | Optic | Nonsocial | 5014758 |
| Fuscatus_2018_81B | Early | Nonsocial | Brain | Nonsocial | 3155963 |
| Fuscatus_2018_85A | Early | Social | Optic | NA_low aggression in trial | 5901566 |
| Fuscatus_2018_85B | Early | Social | Brain | NA_low aggression in trial | 5488583 |
| Fuscatus_2018_87B | Early | Nonsocial | Brain | Nonsocial | 5899623 |
| Fuscatus_2018_88A | Early | Social | Optic | Loser | 9147177 |
| Fuscatus_2018_88B | Early | Social | Brain | Loser | 4059612 |
| Fuscatus_2018_90A | Late | Social | Optic | NA_low aggression in trial | 4375182 |
| Fuscatus_2018_90B | Late | Social | Brain | NA_low aggression in trial | 5571363 |
| Fuscatus_2018_93B | Early | Nonsocial | Optic | Nonsocial | 4143598 |
| Fuscatus_2018_95A | Late | Social | Optic | NA_low aggression in trial | 3335938 |

### Table S2

| Models | BRAIN | | | OPTIC | | |
|---|---|---|---|---|---|---|
| | Up | Down | total | Up | Down | total |
| **Social+Time** | | | | | | |
| Social | 147 | 201 | 348 | 0 | 0 | 0 |
| Time | 5 | 7 | 12 | 0 | 5 | 5 |
| **timeSocial** | | | | | | |
| Early Social - Early Nonsocial | 70 | 137 | 207 | 1 | 0 | 1 |
| Early Social - Late Social | 3 | 0 | 3 | 0 | 0 | 0 |
| Late Social - Late Nonsocial | 0 | 0 | 0 | 0 | 3 | 3 |
| Early Nonsocial - Late Nonsocial | 0 | 0 | 0 | 1 | 1 | 2 |
| Late Social - Early Nonsocial | 76 | 37 | 113 | 1 | 1 | 2 |
| Early Social - Late Nonsocial | 3 | 1 | 4 | 1 | 1 | 2 |
| **Outcome+Tissue+Time** | | | | | | |
| Time | 1 | 8 | 9 | 0 | 0 | 0 |
| W-N | 55 | 42 | 97 | 0 | 0 | 0 |
| L-N | 11 | 2 | 13 | 0 | 0 | 0 |
| W-L | 0 | 0 | 0 | 0 | 5 | 5 |

### Table S3

| GO.ID | Term | Annot. | Sig. | Exp. | P value |
|---|---|---|---|---|---|
| GO:0050909 | sensory perception of taste | 25 | 12 | 2.45E+00 | 1.10E-06 |
| GO:0070588 | calcium ion transmembrane transport | 36 | 13 | 3.53E+00 | 1.80E-05 |
| GO:0045433 | male courtship behavior, veined wing gen... | 27 | 11 | 2.65 | 2.20E-05 |
| GO:0007615 | anesthesia-resistant memory | 38 | 13 | 3.72 | 3.60E-05 |
| GO:0009312 | oligosaccharide biosynthetic process | 10 | 6 | 0.98 | 0.00013 |
| GO:0008039 | synaptic target recognition | 54 | 14 | 5.29 | 0.00029 |
| GO:0030534 | adult behavior | 261 | 46 | 25.57 | 0.00036 |
| GO:0034765 | regulation of ion transmembrane transpor... | 38 | 10 | 3.72 | 0.00043 |
| GO:0001508 | action potential | 12 | 6 | 1.18 | 0.00047 |
| GO:0019722 | calcium-mediated signaling | 31 | 10 | 3.04 | 0.0005 |
| GO:0040034 | regulation of development, heterochronic | 21 | 8 | 2.06 | 0.00052 |
| GO:0016319 | mushroom body development | 107 | 22 | 10.48 | 0.00055 |
| GO:0008345 | larval locomotory behavior | 56 | 14 | 5.49 | 0.00076 |
| GO:0048790 | maintenance of presynaptic active zone s... | 13 | 6 | 1.27 | 0.00081 |
| GO:0048864 | stem cell development | 37 | 5 | 3.63 | 0.00094 |
| GO:0046928 | regulation of neurotransmitter secretion | 28 | 9 | 2.74 | 0.00098 |
| GO:0016079 | synaptic vesicle exocytosis | 51 | 13 | 5 | 0.00117 |
| GO:0007317 | regulation of pole plasm oskar mRNA loca... | 46 | 12 | 4.51 | 0.00119 |
| GO:0044719 | regulation of imaginal disc-derived wing... | 35 | 10 | 3.43 | 0.00143 |
| GO:0030536 | larval feeding behavior | 19 | 7 | 1.86 | 0.00147 |
| GO:0001941 | postsynaptic membrane organization | 10 | 5 | 0.98 | 0.00147 |
| GO:0007271 | synaptic transmission, cholinergic | 10 | 5 | 0.98 | 0.00147 |
| GO:0061028 | establishment of endothelial barrier | 10 | 5 | 0.98 | 0.00147 |
| GO:0007319 | negative regulation of oskar mRNA transl... | 10 | 5 | 0.98 | 0.00147 |
| GO:2000331 | regulation of terminal button organizati... | 48 | 12 | 4.7 | 0.00178 |
| GO:0007411 | axon guidance | 436 | 70 | 42.72 | 0.00233 |
| GO:0007623 | circadian rhythm | 226 | 37 | 22.14 | 0.00245 |
| GO:0097090 | presynaptic membrane organization | 11 | 5 | 1.08 | 0.00248 |
| GO:0007303 | cytoplasmic transport, nurse cell to ooc... | 16 | 6 | 1.57 | 0.00292 |
| GO:0050770 | regulation of axonogenesis | 112 | 21 | 10.97 | 0.0033 |
| GO:0060537 | muscle tissue development | 58 | 13 | 5.68 | 0.00338 |
| GO:0042332 | gravitaxis | 33 | 9 | 3.23 | 0.00349 |
| GO:0007274 | neuromuscular synaptic transmission | 113 | 24 | 11.07 | 0.00354 |
| GO:0043087 | regulation of GTPase activity | 121 | 24 | 11.85 | 0.00377 |
| GO:0048813 | dendrite morphogenesis | 255 | 44 | 24.98 | 0.00388 |
| GO:1900242 | regulation of synaptic vesicle endocytos... | 12 | 5 | 1.18 | 0.00391 |
| GO:0042044 | fluid transport | 12 | 5 | 1.18 | 0.00391 |

| | | | | | |
|---|---|---|---|---|---|
| GO:0016332 | establishment or maintenance of polarity... | 12 | 5 | 1.18 | 0.00391 |
| GO:0009953 | dorsal/ventral pattern formation | 207 | 32 | 20.28 | 0.00393 |
| GO:0035556 | intracellular signal transduction | 596 | 86 | 58.39 | 0.0041 |
| GO:0048808 | male genitalia morphogenesis | 37 | 10 | 3.63 | 0.00452 |
| GO:0007631 | feeding behavior | 85 | 22 | 8.33 | 0.00459 |
| GO:0008016 | regulation of heart contraction | 39 | 10 | 3.82 | 0.00497 |
| GO:0007602 | phototransduction | 86 | 14 | 8.43 | 0.00501 |
| GO:0048167 | regulation of synaptic plasticity | 41 | 10 | 4.02 | 0.00511 |
| GO:0040018 | positive regulation of multicellular org... | 35 | 9 | 3.43 | 0.00535 |
| GO:0007156 | homophilic cell adhesion via plasma memb... | 29 | 8 | 2.84 | 0.00539 |
| GO:0051302 | regulation of cell division | 38 | 8 | 3.72 | 0.00571 |
| GO:0060079 | excitatory postsynaptic potential | 18 | 6 | 1.76 | 0.00571 |
| GO:1900073 | regulation of neuromuscular synaptic tra... | 54 | 11 | 5.29 | 0.00668 |
| GO:0000578 | embryonic axis specification | 180 | 29 | 17.64 | 0.00725 |
| GO:0042220 | response to cocaine | 21 | 7 | 2.06 | 0.00759 |
| GO:0007297 | ovarian follicle cell migration | 193 | 30 | 18.91 | 0.00766 |
| GO:1902600 | proton transmembrane transport | 23 | 5 | 2.25 | 0.0077 |
| GO:0043297 | apical junction assembly | 75 | 17 | 7.35 | 0.00773 |
| GO:0008277 | regulation of G protein-coupled receptor... | 49 | 12 | 4.8 | 0.00784 |
| GO:0051058 | negative regulation of small GTPase medi... | 38 | 7 | 3.72 | 0.00801 |
| GO:0060857 | establishment of glial blood-brain barri... | 25 | 7 | 2.45 | 0.00834 |
| GO:0071907 | determination of digestive tract left/ri... | 14 | 5 | 1.37 | 0.00838 |
| GO:0007616 | long-term memory | 87 | 16 | 8.52 | 0.0094 |
| GO:0019220 | regulation of phosphate metabolic proces... | 294 | 23 | 28.8 | 0.00988 |
| GO:0045727 | positive regulation of translation | 32 | 8 | 3.14 | 0.01016 |
| GO:0007416 | synapse assembly | 288 | 49 | 28.22 | 0.01018 |
| GO:0008049 | male courtship behavior | 89 | 23 | 8.72 | 0.01073 |
| GO:0044089 | positive regulation of cellular componen... | 189 | 30 | 18.52 | 0.01115 |
| GO:0030865 | cortical cytoskeleton organization | 87 | 18 | 8.52 | 0.01132 |
| GO:0046660 | female sex differentiation | 38 | 10 | 3.72 | 0.01157 |
| GO:0051090 | regulation of DNA-binding transcription ... | 28 | 6 | 2.74 | 0.01181 |
| GO:0045450 | bicoid mRNA localization | 10 | 4 | 0.98 | 0.01183 |
| GO:0032368 | regulation of lipid transport | 10 | 4 | 0.98 | 0.01183 |
| GO:0043113 | receptor clustering | 10 | 4 | 0.98 | 0.01183 |
| GO:0006020 | inositol metabolic process | 10 | 4 | 0.98 | 0.01183 |
| GO:0060402 | calcium ion transport into cytosol | 10 | 4 | 0.98 | 0.01183 |
| GO:0071456 | cellular response to hypoxia | 33 | 8 | 3.23 | 0.01229 |
| GO:0007611 | learning or memory | 271 | 53 | 26.55 | 0.01255 |
